# Structural basis of 3′-tRNA maturation by the human mitochondrial RNAse Z complex

**DOI:** 10.1101/2024.05.14.593974

**Authors:** Genís Valentín Gesé, B. Martin Hällberg

## Abstract

Human mitochondrial tRNA maturation is essential for cellular energy production, yet the underlying mechanisms remain only partially understood. Here, we present several cryo-EM structures of the mitochondrial RNase Z complex (ELAC2/SDR5C1/TRMT10C) bound to different maturation states of mitochondrial tRNA^His^, showing the molecular basis for tRNA-substrate selection and catalysis. Our structural insights provide a molecular rationale for the 5′-to-3′ tRNA processing order in mitochondria, the 3′-CCA antideterminant effect, and the basis for sequence-independent recognition of mitochondrial tRNA substrates. Furthermore, the study links mutations in ELAC2 to clinically relevant mitochondrial diseases, offering a deeper understanding of the molecular defects contributing to these conditions.

## Introduction

All human RNA transcripts must be enzymatically processed in different ways to fulfill their biological functions. In the nucleus, RNA polymerase III transcribes tRNA precursors containing 5′- and 3′-leader sequences, which are cleaved by the RNAse P complex and RNAse Z, respectively, to generate tRNAs and other regulatory non-coding RNAs (Altman & Kirsebom, 1999; Siira et al., 2018).

In contrast to nuclear tRNAs, mitochondrial tRNAs are transcribed within two polycistronic heavy-and light-strand precursor RNA transcripts, which contain the coding sequences for mRNAs, rRNAs, and tRNAs. Most RNA moieties in the polycistronic transcripts are separated by tRNA sequences, which serve as landmarks to direct the endonucleolytic cleavage of the polycistronic transcripts by the mitochondrial RNAse P and RNAse Z complexes (punctate model; (Ojala et al., 1981).

Nuclear and mitochondrial RNAse P are evolutionarily and architecturally distinct in humans (Holzmann et al., 2008; Hällberg & Larsson, 2014). Specifically, human nuclear RNAse P is a large RNA-protein complex in which the RNA part catalyzes RNA processing, whereas human mitochondrial RNAse P is a protein-only enzyme (Bartkiewicz et al., 1989) and acts as a complex consisting of TRMT10C/SDR5C1/PRORP (Holzmann et al., 2008). In this complex, PRORP is the nuclease subunit (Holzmann et al., 2008; Reinhard et al., 2015). TRMT10C is a purine N1-methyltransferase that acts at position 9 of mt-tRNAs using S-adenosylmethionine (SAM) as a co-substrate. SDR5C1 is a moonlighting homotetrameric short-chain dehydrogenase/reductase (Holzmann et al., 2008; Vilardo & Rossmanith, 2015). In humans, the nuclease active site is disordered in PRORP and hence is only active in complex with TRMT10C/SDR5C1 (Reinhard et al., 2015).

After tRNA 5′-processing, 3′-processing by RNAse Z ensues. In all domains of life, RNase Z belongs to the metallo-β-lactamase family of enzymes and comprises long and short forms. In prokaryotes, RNAse Z is present only in the short form, which contains one fully functional metallo-β-lactamase domain and functions as a dimer. The long form, which is present along with the short form in most eukaryotes, results from gene duplication and contains two metallo-β-lactamase domains, the NTD and the CTD, where only the CTD has retained the HxHxDH, the PxKxRN (P-loop) and the AxDx motifs important for the enzymatic activity while the NTD contains an insertion known as the exosite that consists of a compact globular domain formed by an insertion in the β-lactamase domain and lies at the tip of a two-stranded stalk (Schilling et al., 2005). In humans, the short-form homolog is named ELAC1 and localizes in the cytosol and the nucleus (Brzezniak et al., 2011; Takahashi et al., 2008; Rossmanith, 2011). The human long form of RNase Z, ELAC2, has two alternative start codons to produce a nuclear and a mitochondrially-targeted form (Brzezniak et al., 2011; Rossmanith, 2011). Accordingly, while nuclear 3′-tRNA processing can be performed by either ELAC1 or ELAC2, only ELAC2 is translocated into mitochondria.

Mutations in ELAC2 are coupled to many clinically relevant mitochondrial diseases, with patient symptoms ranging from pediatric cardiomyopathy to adult intellectual disability depending on their functional severity (Akawi et al., 2016; Cafournet et al., 2023, 2023; Paucar et al., 2018; Saoura et al., 2019). ELAC2 function is also important for nuclear RNA processing of tRNAs and other non-coding RNAs such as sno-RNAs (Siira et al., 2018). To the best of our knowledge, no clinically relevant mutations in ELAC2 have been reported related explicitly to nuclear RNA processing, presumably because mitochondrial dysfunction dominates the clinical picture and nuclear ELAC1 may compensate for reduced ELAC2-mediated nuclear tRNA processing.

We have previously shown that after 5′-leader cleavage by PRORP, most human mitochondrial tRNA precursors remain bound to TRMT10C/SDR5C1, significantly enhancing the efficiency of the ensuing ELAC2 3′-processing for 17 of the 22 human mitochondrial tRNAs (Reinhard et al., 2017). Remarkably, the tRNA precursors also remain bound to TRMT10C/SDR5C1 after ELAC2 processing and even during further maturation. We, therefore, proposed a model in which the TRMT10C/SDR5C1 complex acts as a tRNA-maturation platform in human mitochondria (Reinhard et al., 2017). In this model, most tRNAs are recognized and bound by the TRMT10C/SDR5C1 complex, presumably co-transcriptionally, and then act as a maturation platform to stabilize the partially degenerate human mitochondrial tRNA pool during the ensuing maturation (Reinhard et al., 2017).

After 5′- and 3′-processing, mitochondrial tRNA maturation continues with the addition of a CCA triplet by the CCA-adding enzyme at the 3′-end (Rossmanith et al.,1995; Hällberg & Larsson, 2014), presumably while still bound to the TRMT10C/SDR5C1 tRNA maturation platform. Interestingly, the 3′-CCA is an RNAse Z antideterminant, which is recognized and not removed by ELAC2 to avoid futile cycling of the tRNAs between the CCA-adding enzyme and RNAse Z (Mohan et al., 1999; Nashimoto, 1997). The 3′-CCA antideterminant effect ensures an efficient tRNA maturation process – the lack of which leads to a spectrum of diseases similar to those observed when CCA addition is impaired (Wedatilake et al., 2016).

Despite the importance of ELAC2 functionality for human health, the molecular basis of tRNA recognition, catalysis, and CCA antidetermination remains unknown. Here, we determined the structures of several ternary complexes of human mitochondrial RNAse Z. Our structures reveal the molecular details of ELAC2’s interactions with its tRNA substrate and the TRMT10C/SDR5C1 tRNA-maturation platform. Furthermore, we show how ELAC2 discriminates against tRNAs with an unprocessed 5′-end or those bearing a 3′-CCA tail and provide a molecular rationale for several clinically relevant mutations.

## Results

### Cryo-EM structure of the mitochondrial RNAse Z product ternary complex

To understand the structural basis of mitochondrial tRNA 3′-processing, we reconstituted the human mitochondrial RNAse Z-complex using recombinantly expressed TRMT10C–SDR5C1 subcomplex, a tRNA precursor, and ELAC2 (Figure 1A; Supplementary Fig. 1A). As a precursor tRNA substrate, we used the tRNA cluster tRNA^His^-tRNA^Ser(AGY)^ (abbreviated as HS) located on the mtDNA heavy strand (Figure 1B). The reconstituted catalytically active RNAse Z (Supplementary Fig. 1B) was incubated to complete turnover and used to determine a cryo-EM structure of the ternary product complex of human RNAse Z to an overall resolution of 2.8 Å (Figure 1C,D; Supplementary Figs. 2 and 3 and Table 1). Within the complex, a tetramer of SDR5C1 forms a flat platform that can be bound from each flat side by TRMT10C and precursor mt-tRNA. Thus, the SDR5C1/TRMT10C subcomplex can support the simultaneous processing of two tRNA molecules. As the SDR5C1 platform has two-fold symmetry within the plane, we observed TRMT10C/tRNA in either of the two equivalent positions rotated by 180° to each other (Supplementary Fig. 4).

**Figure 1.**
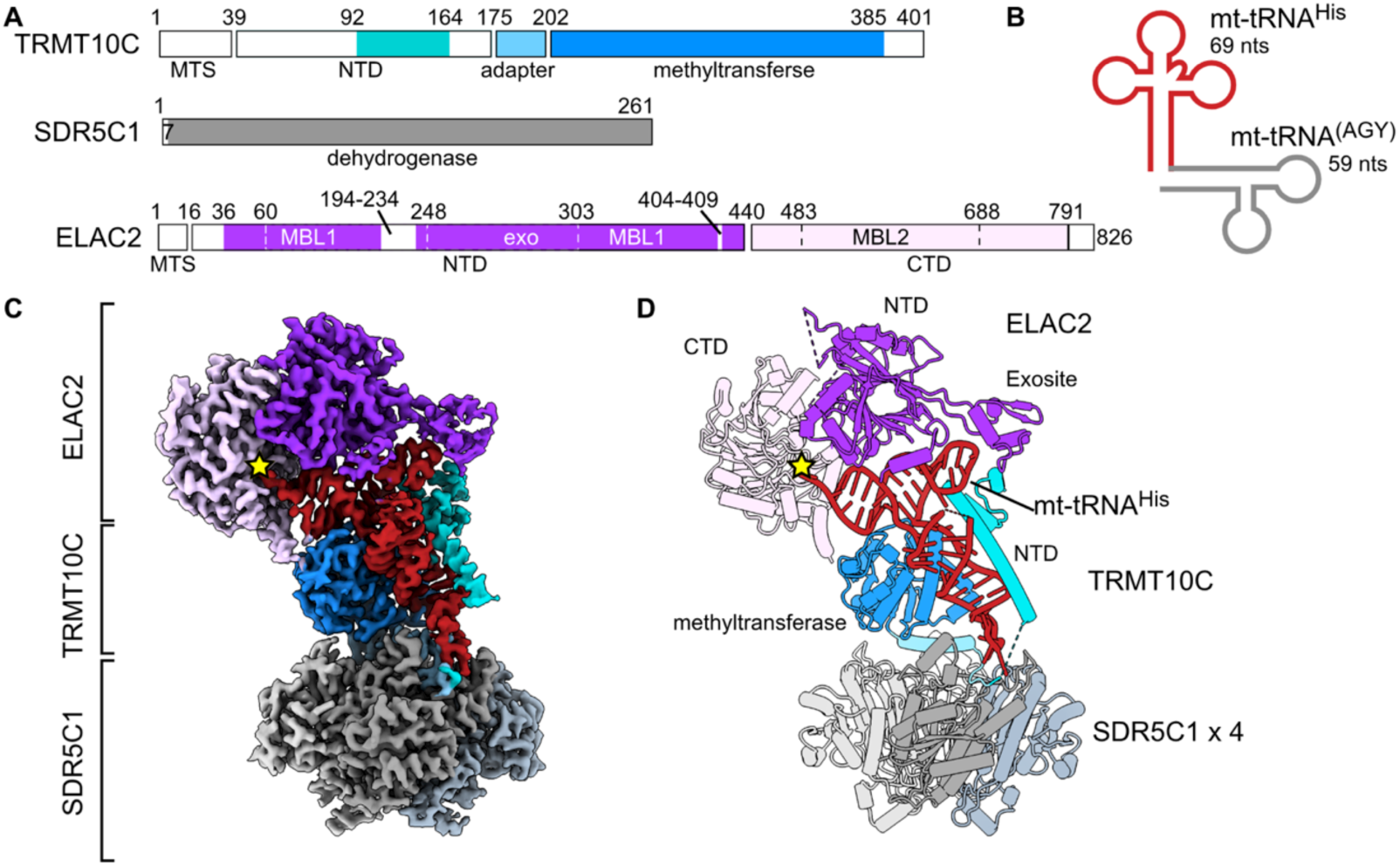
Structure of human mitochondrial RNAse Z. **A.** Domain architecture of the RNAse Z components. The N-terminal (NTD) and C-terminal (CTD) domains of ELAC2 are shown in dark and light purple, respectively. MBL1 and MBL2, which are the metallo-β-lactamase folds, and the exosite insertion are indicated with dotted lines. TRMT10C is shown in blue, with the NTD highlighted in cyan. MTS, mitochondrial targeting sequence. Unmodelled regions are in white. **B.** The mt-tRNA precursor consisted of tRNA^His^ (red) and tRNA^Ser(AGY)^. **C** and **D**. Cryo-EM reconstruction (C) and cartoon representation (D) of the RNAse Z complex. Colored as in A. The four SDR5C1 subunits are shown in different shades of gray. The ELAC2 active site is marked with a yellow star.

**Table 1.** Cryo-EM data collection, refinement, and validation statistics

|  | RNAse Z-HS<br>(EMDB-50050)<br>(PDB 9EY0) | RNAse Z<sup>H548A</sup>-HS<br>(EMDB-50051)<br>(PDB 9EY1) | RNAse Z-HCCA<br>(EMDB-50052)<br>(PDB 9EY2) |
| --- | --- | --- | --- |
| **Data collection and processing** |  |  |  |
| Magnification | 165,000 | 165,000 | 165,000 |
| Voltage (kV), source | 300, XFEG | 300, XFEG | 300, E-CFEG |
| Electron exposure (e<sup>-</sup>/Å<sup>2</sup>) | 59 | 59 | 59 |
| Defocus range (μm) | 0.2-2 | 0.2-2 | 0.2-2 |
| Pixel size (Å) | 0.505 | 0.505 | 0.505 |
| Symmetry imposed | P1 | P1 | P1 |
| Initial particle images (no.) | 293,145 | 643,629 | 4,773,315 |
| Final particle images (no.) | 28,771 | 15,961 | 52,932 |
| Map resolution (Å) | 2.8 | 2.7 | 2.1 |
| FSC threshold | 0.143 | 0.143 | 0.143 |
| Map resolution range (Å) | 2.5-7 | 2.5-7 | 2.2-7 |
| **Refinement** |  |  |  |
| Initial model used (PDB code) | 7ONU | 7ONU | 7ONU |
| Model resolution (Å) | 3.0 | 2.8 | 2.2 |
| FSC threshold | 0.5 | 0.5 | 0.5 |
| Model composition |  |  |  |
| Non-hydrogen atoms | 16,691 | 16,783 | 16,764 |
| Protein/RNA residues | 2008/65 | 2009/69 | 2009/68 |
| Ligands | GTP:1 Zn:2 SAM:1 | GTP:1 Zn:1 SAM:1 | GTP:1 Zn:2 SAM:1 |
| *B* factors (Å<sup>2</sup>) |  |  |  |
| Protein | 22.1/186.4/74.99 | 0/106.2/35.1 | 19.67/222.34/71.33 |
| RNA | 40.5/147.0/90.5 | 9.8/167.9/70.0 | 14.56/169.09/65.64 |
| Ligand | 42.8/189.2/85.3 | 78.55/171.39/91.80 | 94.89/124.68/100.82 |
| R.m.s. deviations |  |  |  |
| Bond lengths (Å) | 0.004 | 0.005 | 0.003 |
| Bond angles (°) | 0.540 | 0.634 | 0.543 |
| Validation |  |  |  |
| MolProbity score | 1.36 | 1.38 | 1.88 |
| Clashscore | 4.49 | 4.90 | 6.78 |
| Poor rotamers (%) | 0.18 | 0.43 | 3.74 |
| Ramachandran plot |  |  |  |
| Favored (%) | 97.29 | 97.34 | 97.69 |
| Allowed (%) | 2.71 | 2.66 | 2.26 |
| Disallowed (%) | 0.00 | 0.05 | 0.00 |

The SDR5C1/TRMT10C/tRNA part of the RNAse Z product complex is similar to a previously published mt-RNAse P structure, in which the SDR5C1/TRMT10C subcomplex interacts with PRORP to catalyze the 5′-cleavage of a precursor mt-tRNA (Bhatta et al., 2021). This highlights the notion that mt-tRNA processing by mt-RNAse P and mt-RNAse Z is tightly coupled (Reinhard et al., 2017) as it is only required to exchange the RNA processing catalytic subunit without major structural rearrangements in the SDR5C1/TRMT10C/tRNA platform. In our product complex, D314–P319 in TRMT10C, known as motif II in the TrmD-family of tRNA methyltransferases, have a different conformation compared to that observed in the apo-TRMT10C structure (Bhatta et al., 2021). Here, the motif II loop’s conformation is similar to the SAM-bound TRMT10C methyltransferase (PDB: 5NFJ), and we observed SAH in the methyltransferase active site (Supplementary Fig. 4).

Furthermore, the structure shows how TRMT10C compensates for the lack of strong D-arm/T-arm interactions in most bilaterian mitochondrial tRNAs by providing structural support through its extensive interactions with the mt-tRNA’s D arm. In addition, TRMT10C’s N-terminal domain stabilizes the T-arm loop via electrostatic interactions with the RNA backbone (Figure 2A–C). Collectively, these interactions underscore TRMT10C’s importance in promoting bilaterian mt-tRNA folding, and these elements are recognized by ELAC2 when in the complex with the SDR5C1/TRMT10C tRNA-processing platform.

**Figure 2.**
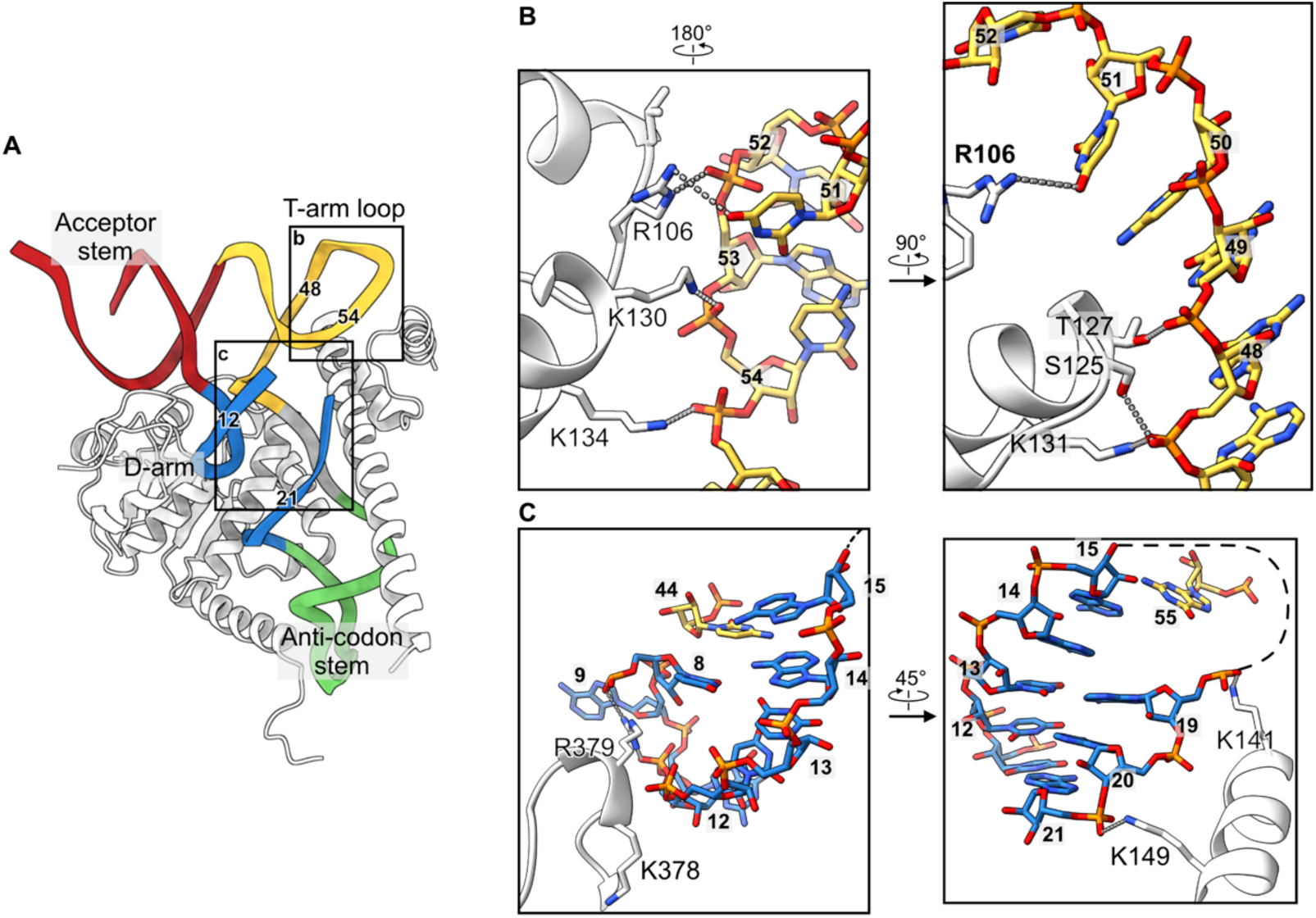
Stabilization of the tRNA fold by TRMT10C. **A.** TRMT10C (white) wraps around the mt-tRNA^His^ anticodon stem and D-arm to stabilize the cloverleaf fold. The nucleotide numbers forming the T-arm loop are indicated for comparison with those in panels B and C. **B** and **C**. Molecular interactions of TRMT10C with the T-arm loop (**B**) and the D-arm (**C**). The views have been rotated for easier visualization.

The structure of human ELAC2 is similar to that of the yeast homolog Trz1. It consists of two β-lactamase domains (N-terminal domain (NTD) and C-terminal domain (CTD)) with insertions and extensions (Figure 1). Although the NTD and CTD have the same fold (RMSD of 2.1 Å for 212 Cα atoms), the NTD lacks the key Zn-bound catalytic site necessary for hydrolytic activity. In addition, similar to Trz1, the human ELAC2 active site on the CTD is situated in a channel formed between the NTD and the CTD (Figure 1C).

In the mitochondrial RNase Z complex, ELAC2 binds to the mt-tRNA T-arm loop and the TRMT10C N-terminal domain through the exosite (Figure 3A,B). The exosite establishes electrostatic interactions with the T-arm loop backbone U52 via K279 and N253. Furthermore, the exosite forms a hydrophobic pocket with F254 at the bottom. This pocket is vacant in our product ternary structure. Still, it might accommodate base stacking on F254 in mt-tRNAs with additional nucleotides in the Ψ-loop, and F254 is conserved in bilaterian ELAC2 homologs (Supplementary Fig. 6A). Furthermore, the exosite interacts with and is stabilized in its position by the TRMT10C N-terminal domain. Here, ELAC2 residues V256, L257, and K260 form hydrophobic interactions with the tip of the TRMT10C N-terminal α-helix, including L103 and L104. The hydrophobic character of these positions is conserved in bilaterians and, therefore, their capacity to form a hydrophobic zipper that links ELAC2 to the SDR5C1/TRMT10C tRNA-processing platform (Supplementary Fig. 6A) to form a clamp over the tRNA T-arm loop (Figure 3B). Notably, the TRMT10C N-terminal α-helix is also strongly contributing to the formation of the mitochondrial RNAse P complex through clamping the tRNA T-arm loop and PRORP (Supplementary Fig. 7A).

**Figure 3.**
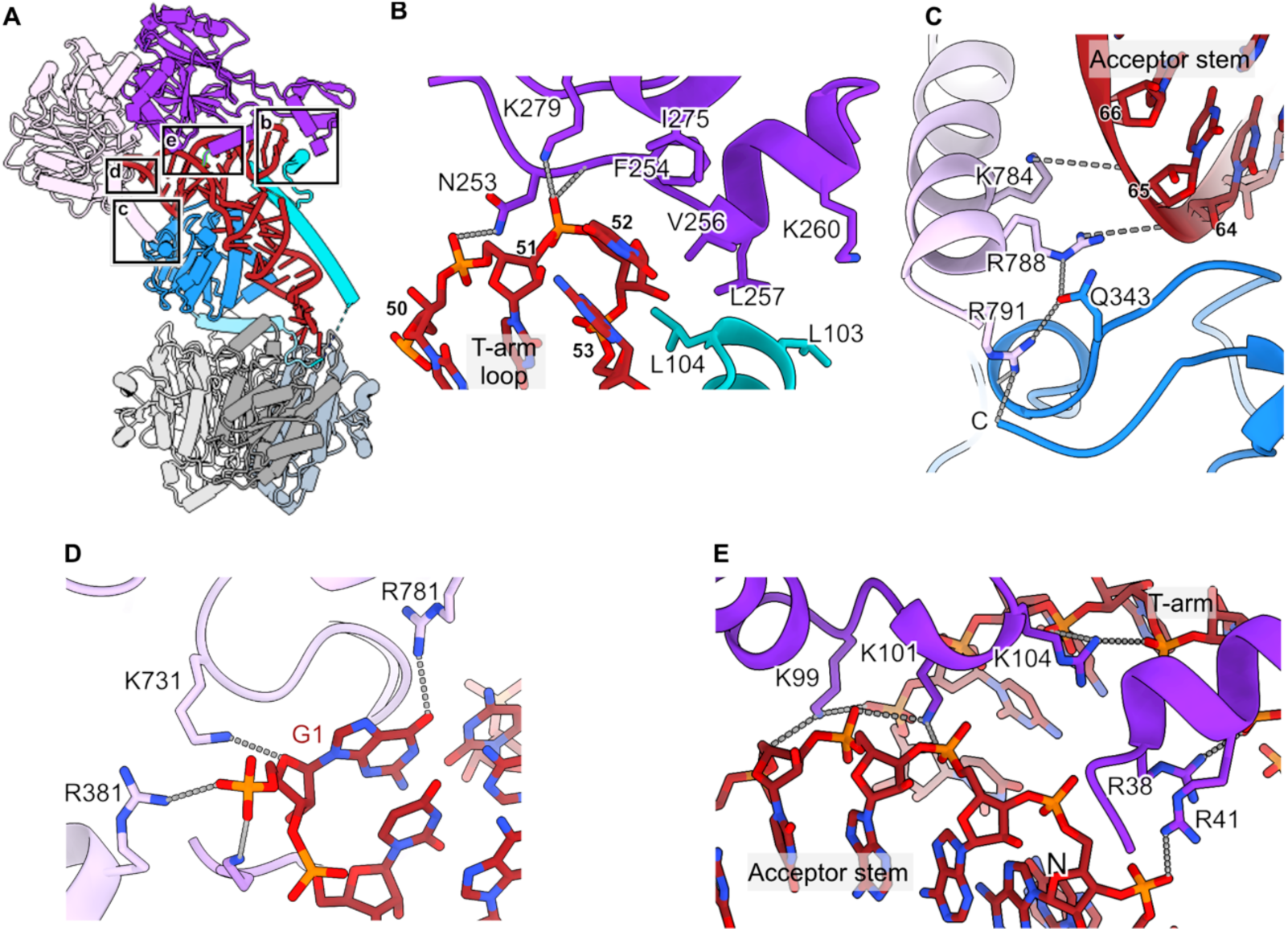
Interaction of ELAC2 with the precursor mt-tRNA^His^. **A**. Overview of the mitochondrial RNAse Z complex. The insets from panels B-E are shown. **B**. Contacts between the ELAC2 exosite, the T-arm loop, and the TRMT10C N-terminal domain. **C**. Contacts between the ELAC2 C-terminal helix, the TRMT10C C-terminus, and the acceptor stem. Small numbers correspond to RNA nucleotides. **D**. Recognition of the 5′-end of the mt-tRNA. Only the α-phosphate of G1 is shown since natural ELAC2 substrates carry a 5′-monophosphate. **E**. Interaction of ELAC2 with the acceptor stem/T-arm groove.

The ELAC2 C-terminal α-helix (α25) contributes further to mitochondrial RNAse Z complex formation (Figure 3C) and the C-terminal α-helix is only present in long-form homologs (Supplementary Fig. 6B). The C-terminal helix contains R788, which interacts with the backbone of U66, located close to the tip of the acceptor stem, as well as with TRMT10C Q343. It also contains R791, which binds to the TRMT10C C-terminal carboxyl group and Q343 (Figure 3C). Notably, this C-terminal α helix would collide with a 5′-extension of the mt-tRNA (Supplementary Fig. 7B) and therefore provides the structural basis for the 5′-to-3′ processing order observed in vivo and in vitro (Rackham et al., 2016; Reinhard et al., 2017). In addition, ELAC2 residues R381, K495, and K731 interact with the 5′-α-phosphate of the mt-tRNA^His^ (Figure 3D). Thus, these three residues may probe if correct 5′-processing has been performed previously by RNAse P through electrostatic interactions with the 5′-monophosphate group.

ELAC2 establishes sequence-independent electrostatic interactions with the major groove between the acceptor stem and the T-arm loop (Figure 3E). The conserved lysines and arginine in the 99-KLKVAR-104 motif (Supplementary Fig. 6C) align well to interact with three consecutive backbone phosphates. Furthermore, two conserved residues, R38 and R41 (Supplementary Fig. 6D), in the ELAC2 N-terminal α helix (α1) line up with and assist the 99-KLKVAR-104 motif in acceptor-arm recognition (Figure 3E). In total, ELAC2 possesses five positively charged side chains that behave like a zipper to anchor the enzyme on the bound mt-tRNA.

Taken together, these results show that ELAC2 recognizes the basic structural features of mt-tRNAs through indirect readout. In addition, ELAC2 is held in place and straddled over the tRNA 3′-end by its N- and C-terminal interactions with TRMT10C on the TRMT10C/SDR5C1 tRNA-maturation platform (Figure 3).

### Structural basis of human mitochondrial RNase Z catalysis

In our product complex RNAse Z-HS, the discriminator base C69 is positioned in the active site, with its ribose-O3 in the expected position directly after catalysis (Figure 4A). Furthermore, the C69 backbone phosphate is wedged between S490, S768, and K700, whereas R728 in the conserved 724-HFSQRY-729 motif (Supplementary Fig. 6E) clamps the C69 and C68 backbone phosphates (Figure 4A). Several clinically relevant mutations associated with hypertrophic cardiomyopathy have been identified in human patients. In particular, three of them affect the substrate-binding residues in and surrounding the HFSQRY-motif and reduce the k_cat_/K_M_ of ELAC2 (Saoura et al., 2019). A clinically relevant mutation in the HFSQRY-motif is Y729C. Here, Y729 stacks on R728 and steers R728 towards the precursor 3′-end (Figure 4A). Another clinically relevant mutation in relation to the HFSQRY-motif is P493L. Here, Q727 in the HFSQRY-motif is strongly held (2.2 Å H-bond) in place by the backbone carbonyl of I492. A P493L mutation will significantly affect this interaction, thus preventing Q727 from remaining anchored as needed to ensure that R728 is properly aligned (Figure 4A). The third clinically relevant mutation related to the HFSQRY-motif is R781H. Here, R781 binds and fixates the R728 backbone carbonyl, which is crucial for R728 to be properly oriented (Figure 4A).

**Figure 4.**
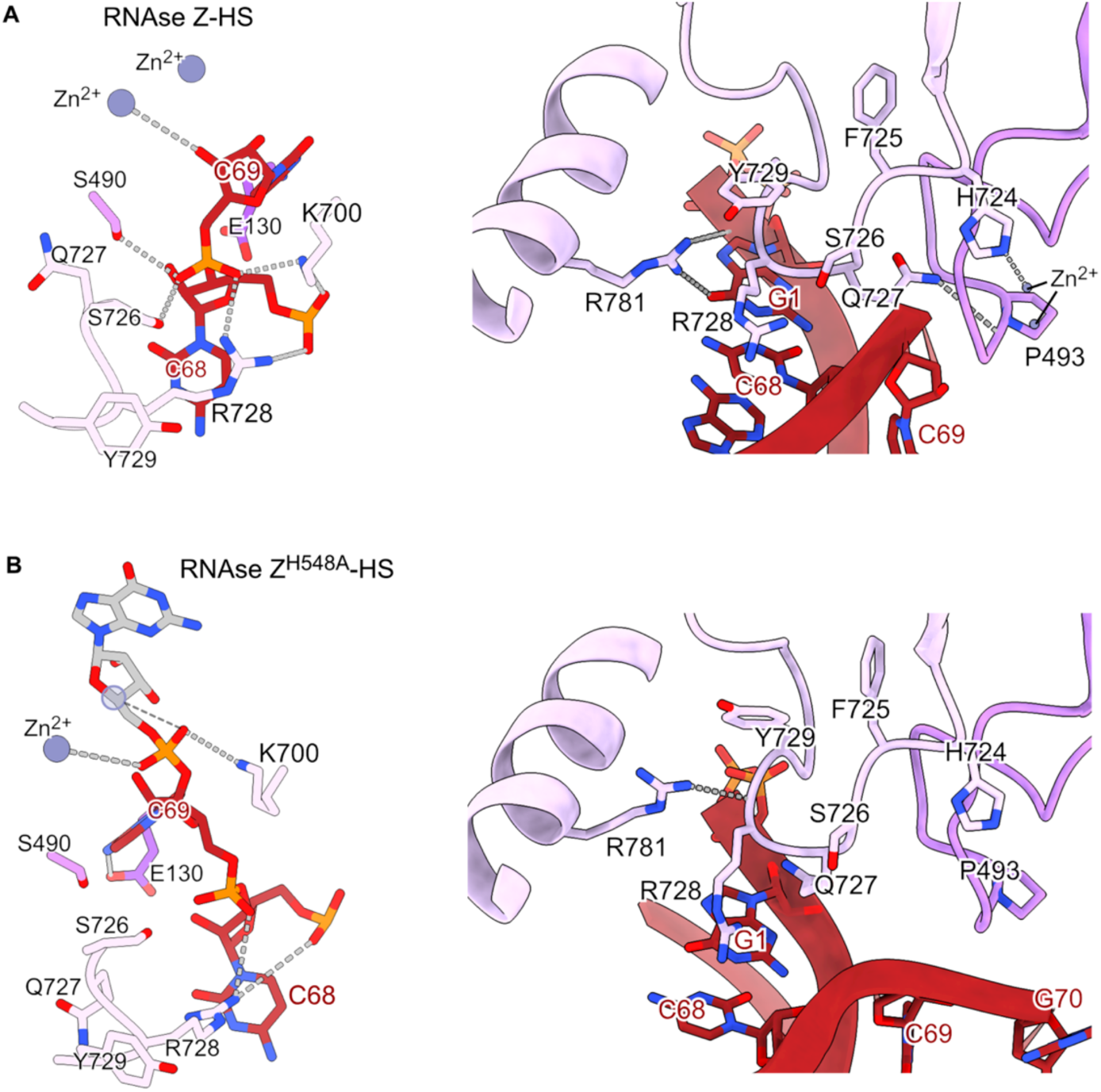
Interactions around the tRNA scissile bond and the ELAC2 HFSQRY motif. **A.** In the RNAse Z-HS structure, the 3′-oxygen interacts with a Zn^2+^ after the 3′-trailer is removed (left panel). **B.** The RNAse Z^H548A^-HS structure shows how the scissile bond is positioned for catalysis (left panel). The missing Zn^2+^ due to the H548A mutation is shown by a transparent circle based on the RNAse Z-HS structure.

To trap an mt-RNAse Z substrate-relevant complex, we mutated H548, which coordinates one of the two active-site Zn^2+^ ions and determined a 2.7 Å resolution cryo-EM structure from a mitochondrial RNAse Z (H548A) complex preparation in which the mt-tRNA precursor was predominantly uncleaved (RNAse Z^H548A^-HS; Supplementary Fig. 1B; Supplementary Figs. 8 and 9 and Table 1). Overall, the structure of the TRMT10C/SDR5C1/mt-tRNA platform did not change with the introduction of the H548A mutation. However, ELAC2 pivots on the mt-tRNA by approximately 3.5° (Supplementary Fig. 10) compared with its position in the product complex. Due to this pivoting, the RNAse Z^H548A^-HS structure shows a difference in the interactions around the discriminator nucleotide and the scissile bond (Figure 4B) compared with the product complex. In the RNAse Z^H548A^-HS structure, the 3′-oxygen of the scissile bond is properly oriented for hydrolysis, but it is displaced by approximately 3 Å, and the discriminator nucleotide C69 moves closer to E130. This may be attributed to the absence of the first Zn^2+^ ion due to the H548A mutation. Notably, K700 switches from interacting with C68 in the product complex RNAse Z-HS to interacting with the scissile bond, *i.e.*, the C69-G1 phosphodiester bond in the RNAse Z^H548A^-HS structure (Figure 4B).

The scissile bond is followed by the mt-tRNA^Ser^ acceptor stem. Only the first nucleotide (G1) was modeled; nevertheless, the cryo-EM map reveals that the acceptor stem contacts ELAC2 residues G156-P157 (Supplementary Fig. 11), which are situated on a ridge in the 3′-direction of the substrate RNA. Consequently, the peptide backbone undergoes a conformational change. This suggests that G156 and P157 are involved in substrate binding, and this is further supported by the clinically relevant F154L mutation since F154 plays a structural role in the stabilization of this ridge (Saoura et al., 2019).

Our RNAse Z^H548A^-HS structure is comparable to the X-ray structure of *B. subtilis* RNAse Z obtained using a non-cleavable substrate analog (Pellegrini et al., 2012). The human ELAC2 G1 corresponds to U1 in *B. subtilis*. The U1 inserts into a pocket between the two β-lactamase domains, and this interaction is necessary for hydrolysis. In human ELAC2, G1 is sterically occluded from the pocket by L126. This indicates that, unlike in *B. subtilis*, the pocket is not used to process the mt-tRNA^His-Ser(AGY)^ precursor (Supplementary Fig. 12).

Taken together, in comparison with the RNAse Z-HS product structure, the RNAse Z^H548A^-HS structure reveals additional ELAC2 residues involved in binding to the mt-tRNA precursor and in orienting the scissile bond, namely E130, K700, and G156-P157.

### Structural basis for the 3′-CCA tail as an antideterminant for ELAC2 processing

After maturation of the mt-tRNA ends by RNAse P and RNAse Z, the mitochondrial CCA-adding enzyme adds the 3′-terminal cytosine-cytosine-adenine (3′-CCA) triplet. 3′-CCA is necessary for the aminoacylation of tRNAs, and thus, it should be protected from ELAC2 activity to prevent futile cycles of addition and removal. Indeed, ELAC2 activity on cytosolic tRNAs bearing the 3′-CCA tail is very inefficient (Mohan et al., 1999; Nashimoto, 1997).

To evaluate the antideterminant effect of the 3′-CCA tail in the mitochondria, mt-tRNA^His^ carrying a 3′-CCA tail was incubated with TRMT10C/SDR5C1 and increasing concentrations of ELAC2. The removal of the 3′-CCA tail by ELAC2 was very inefficient compared to the cleavage of mt-tRNA^His-Ser(AGY)^ (Figure 5A).

**Figure 5.**
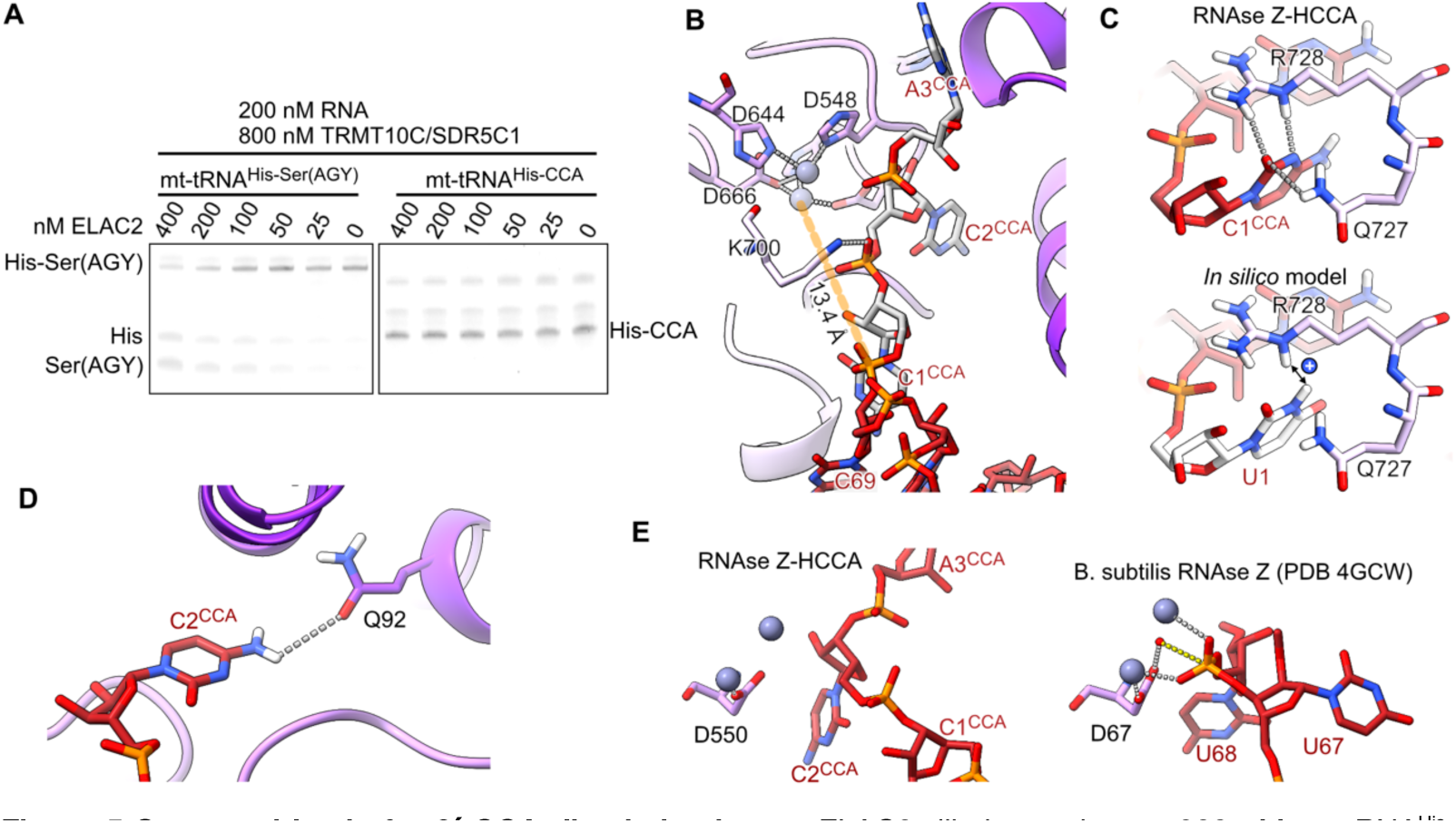
Structural basis for 3’-CCA discrimination. **a**. ELAC2 dilution series on 200 nM mt-tRNA^His-Ser(AGY)^ or mt-tRNA^His-CCA^ and 800 nM TRMT10C/SDR5C1. **b**. The 3’-CCA (gray) occupies the active site channel of ELAC2. The mt-tRNA^His^ (red) 3′-end is 13.4 Å away from the catalytic Zn^2+^ (gray spheres). **c**. Interactions of C1^CCA^ (upper) with ELAC2 R728 and Q727. Replacing C1 with uracil *in silico* (lower) would presumably cause electrostatic repulsion with R728 (blue circle with +). **d**. Interaction of C2^CCA^ with Q92. The hydrogen atoms were added in ChimeraX to visualize the H-bonding interactions. **e**. Comparison with the tRNA^Thr^ precursor bound to the *B. subtilis* RNAse Z (Pellegrini et al., 2012), which shows the interactions required for catalysis. The phosphate oxygens coordinate one Zn^2+^ ion each. The hydrolyzing water molecule (red sphere) is positioned by D550. In RNAse Z-HCCA, the backbone adopts a different conformation, which is unsuitable for catalysis.

To understand the molecular details of 3′-CCA discrimination, we determined the cryo-EM structure to 2.1 Å resolution of RNAse Z with mt-tRNA^His^ carrying the 3′ -CCA tail (RNAse Z-HCCA, Supplementary Figs. 13 and 14 and Table 1).

The TRMT10C/SDR5C1 platform in the RNAse Z-HCCA complex is similar to the RNAse Z-HS and RNAse Z^H548A^-HS complexes, but here, ELAC2 pivots by approximately 6 degrees around the tRNA (Supplementary Fig. 10). Consequently, the active site is now ∼13 Å away from the mt-tRNA^His^ 3′-end, and the active site channel is occupied by the 3′-CCA tail (Figure 5B).

The first C (C1^CCA^) of the 3′-CCA tail is recognized by H-bonding to Q727 and R728 (Figure 5C), which lie at the entrance of the active site channel. The second C (C2^CCA^) inserts into a pocket in the NTD situated on the opposite side of the active site channel. C2^CCA^ establishes H-bonds with Q92, which pulls C2^CCA^ toward the bottom of the pocket (Figure 5D). Furthermore, the phosphodiester bond between C1^CCA^ and C2^CCA^ is stabilized by K700 (Figure 5B). The terminal A (A3^CCA^) base in the CCA does not establish H-bonds. It, therefore, does not seem to participate in the antideterminant effect in accordance with previous in vitro work (Nashimoto, 1997).

The C1^CCA^ – C2^CCA^ bond is the closest to the active site. Nevertheless, it is not well positioned for hydrolysis since it is 1.9 Å further away from the active site than the scissile bond in the mt-tRNA^His^-tRNA^Ser(AGY)^ complex. Furthermore, the phosphate oxygens point away from the active site Zn^2+^, so they cannot participate in the hydrolysis reaction catalyzed by ELAC2 (Figure 5E). This conformation of the RNA backbone is enforced by the interactions established by both C1 and C2.

## Discussion

The structural analysis presented here reveals how ELAC2 interacts with the tRNA processing platform formed by TRMT10C/SDR5C1 to form human mitochondrial RNAse Z.

The binding of ELAC2 through both its N-terminal and C-terminal domains to TRMT10C underscores a complex and finely tuned interaction network essential for the precise positioning and enzymatic activity of ELAC2 on its tRNA substrate. ELAC2 straddling the tRNA molecule, facilitated by its interactions at both terminal domains, exemplifies an indirect readout mechanism in which the enzyme recognizes and interacts with the structural features of tRNA rather than directly binding to specific nucleotide sequences. This mode of recognition allows ELAC2 to effectively process a variety of tRNA sequences by accommodating variations within the tRNA structure, thus maintaining a high degree of processing fidelity across different tRNA species. The dual interaction sites on ELAC2 enhance its stability and positioning on the tRNA-bound TRMT10C/SDR5C1 platform, aligning ELAC2 optimally for catalysis. This structural arrangement is crucial for the enzymatic function of ELAC2, as it ensures that the enzyme is correctly oriented relative to the tRNA substrate. The N-terminal interaction likely contributes to the initial recognition and binding of the tRNA, positioning ELAC2 to engage with the substrate. Subsequently, the C-terminal interaction stabilizes the complex during the catalytic process, ensuring efficient and precise cleavage at the 3′-end of the tRNA. Moreover, the strategic positioning of ELAC2 over the tRNA, secured by interactions at both terminals, indicates a robust regulatory mechanism. This setup facilitates the catalytic action required for tRNA maturation and prevents aberrant processing that could lead to dysfunctional tRNA molecules. Thus, the interaction between ELAC2 and the tRNA maturation platform reflects a highly evolved and specialized system designed to meet the unique demands of mitochondrial biology, where precision in tRNA processing is directly linked to the organelle’s efficiency in protein synthesis.

We have previously shown that the TRMT10C/SDR5C1 complex enhances ELAC2 activity on most mt-tRNA precursors, with the exceptions of mt-tRNA^Gln^, mt-tRNA^Asn^, mt-tRNA^Leu(UUR)^, and mt-tRNA^Ser(UCN)^ (Reinhard et al., 2017). mt-tRNA^Ser(UCN)^ belongs to the Type I tRNAs, which has an extra base pair in the anticodon stem and is not methylated at position 9 (Suzuki et al., 2020; Watanabe, 2010), suggesting that it does not bind to TRMT10C/SDR5C1. Mitochondrial tRNA^Leu(UUR)^, tRNA^Asn,^ and tRNA^Gln^ belong to Type 0 (canonical) tRNAs and are the only mt-tRNAs that contain dihydrouridine and pseudouridine motifs in the D and T loops, respectively (Helm et al., 2000; Suzuki et al., 2020). Also belonging to Type 0 are cytosolic tRNAs, which are processed by ELAC2 in the nucleus (Siira et al., 2018) without TRMT10C/SDR5C1. In contrast, the other mt-tRNAs, which are processed more efficiently in the presence of TRMT10C/SDR5C1, belong to type II tRNAs (Watanabe, 2010), which lack the D loop-T loop interaction (Helm et al., 2000; Wakita et al., 1994). Consequently, the folding of the T loop is less stable in type II tRNAs. Here, the structures show that the NTD of TRMT10C binds to the tRNA T-loop to stabilize its fold and facilitate the tRNA interaction with the ELAC2 exosite. Furthermore, the TRMT10C/SDR5C1 tRNA-maturation platform’s role extends beyond mere structural support; it also likely impacts the kinetic properties of the processing enzymes. By pre-organizing the tRNA substrates, the platform may reduce the entropy cost during the transition state of the enzymatic reaction, thus enhancing the catalytic efficiency. This pre-organization is crucial in mitochondria, where the compact genome and the need for rapid and accurate protein synthesis necessitate highly efficient processing mechanisms.

The observations above underscore the role of TRMT10C/SDR5C1 as a platform for mt-tRNA processing, and this is further supported by two preprints made available during the preparation of this manuscript (Bhatta et al., 2024; Meynier et al., 2023). As such, it supports the multistep processing of mt-tRNA precursors from 5′-endonucleolytic cleavage by PRORP, 3′-endonucleolytic cleavage by ELAC2, and 3′-CCA addition. These steps occur in strict sequential order (Lopez Sanchez et al., 2011; Rackham et al., 2016; Reinhard et al., 2017; Rossmanith et al., 1995). Our structure of RNAse Z shows that the ELAC2 C-terminal helix (α24) would sterically clash with 5′-unprocessed pre-tRNAs (Supplementary Fig. 7B) and, therefore, forcing ELAC2 to act after RNAse P. In addition, ELAC2 interactions with the 5′-phosphate via R381, K495, and K731 are important for positioning ELAC2 for catalysis.

In our product complex RNase Z-HS structure, the discriminator base C69 is held in place by interactions with S490, S768, and K700, while R728 from the 724-HFSQRY-729 motif secures the backbone phosphates of C69 and C68. Notably, mutations within or structurally adjacent to the HFSQRY motif, such as Y729C, P493L, and R781H, are clinically significant (Saoura et al., 2019). These mutations disrupt substrate binding and enzymatic efficiency, as evidenced by their impact on the orientation and stability of key residues necessary for tRNA processing. For example, Y729C affects the positioning of R728, which is critical for interacting with the tRNA precursor’s 3′-end, whereas P493L compromises the anchoring of Q727, affecting the alignment necessary for catalytic activity. Similarly, R781H disrupts the positioning of R728, illustrating how mutations in the HFSQRY motif can have profound effects on the function of ELAC2, with implications for diseases such as hypertrophic cardiomyopathy.

After 3′-cleavage by ELAC2, the 3′-CCA tail is added to mt-tRNA precursors, which is required for aminoacylation. To prevent futile cycles of 3′-CCA tail addition and removal, ELAC2 recognizes and ignores precursor tRNAs that carry the 3′-CCA tail (Nashimoto, 1997). Our RNAse Z-HCCA structure shows that ELAC2 recognizes the 3′-CCA tail by interacting with the first two “C” nucleotides: C1^CCA^ and C2^CCA^. Specifically, C1^CCA^ is recognized by H-bonding to Q727 and R728 in the sequence motif 724-HFSQRY-729 (Supplementary Fig. 6E), where R728 is conserved, and Q727 is conserved in long-form RNase Z homologs. Based on our structure, purines are likely sterically occluded from the C1^CCA^-binding site, whereas uracil, on the other hand, is disfavored due to a mismatching H-bond pattern (Figure 5D). Indeed, the mutation of uracil to cytosine in mt-tRNA^Ser(UCN)^ generates a 3′-CCU trailer that cannot be processed by mitochondrial RNAse Z and is a cause of non-syndromic deafness (Levinger et al., 2001). Furthermore. C2^CCA^ inserts into a pocket in the NTD of ELAC2, where it interacts with Q92. This interaction pulls the C1-C2 phosphodiester bond away from the active site, thereby preventing hydrolysis.

The role of the HFSQRY motif in ELAC2 presents a fascinating example of multifunctionality within mitochondrial tRNA processing enzymes. This motif is pivotal not only for the catalytic activity of ELAC2 but also plays a crucial role in recognizing the CCA end of tRNAs, which act as an antideterminant for ELAC2 catalysis. Therefore, mutations in the HFSQRY motif can impact both the catalytic activity of ELAC2 and its ability to recognize the CCA sequence correctly, and it is tempting to speculate that the clinically relevant mutations that have been identified in this motif may have an intermixed, double effect on mitochondrial 3′-tRNA maturation.

## Methods

### Cloning and protein expression

The expression constructs for TRMT10C/SDR5C1 and ELAC2 were as previously described (Reinhard et al., 2017). The H548A mutation was introduced by site-directed mutagenesis (using Pfu Ultra II from Agilent Technologies). Expression was performed in *E. coli* KRX cells (Promega) in phosphate-buffered Terrific Broth (Sigma-Aldrich) supplemented with 50 μg/ml kanamycin. Expression was induced at OD_600_ 0.8 with 0.5 mM isopropyl β-D-1-thiogalactopyranoside (IPTG) and rhamnose overnight at 18 °C.

### Protein purification

Cells were lysed in buffer A (50 mM HEPES pH 7.5, 300 mM NaCl, 1 mM β-mercaptoethanol, 5% glycerol, 1 mM IPTG, DNAse I (Grade I, Roche)). Ni^2+^ IMAC chromatography (Cytiva) was run with buffer A and eluted with buffer B (20 mM HEPES pH 7.5, 150 mM NaCl, 1 mM β-mercaptoethanol, 5% glycerol, 300 mM imidazole). Next, the sample was directly applied and run through a HiTrap Q column (Cytiva) pre-equilibrated with buffer C (20 mM HEPES pH 7.5, 150 mM NaCl, 5% glycerol, 1 mM DTT) to remove contaminating nucleic acids. The sample was applied on a HiTrap HEPARIN (Cytiva) pre-equilibrated with buffer C and eluted with a 1 M NaCl gradient. The sample was dialyzed overnight in buffer C at 4 °C in the presence of TEV protease (in-house production) and run on a Capto S column (Cytiva) eluted with a 1 M NaCl gradient. Finally, TRMT10C/SDR5C1 was run through a Sephacryl S300 (Cytiva) in buffer C. TRMT10C/SDR5C1 was frozen in liquid nitrogen and stored for later use.

ELAC2 was lysed and purified using Ni^2+^ IMAC chromatography as above. Next, ELAC2 was dialyzed in buffer C in the presence of TEV protease. ELAC2 was run through a HiTrapQ column (Cytiva) in buffer C to remove contaminating nucleic acids and purified over a HiTrap HEPARIN column in buffer C with a salt gradient up to 1 M NaCl. ELAC2 was frozen in liquid nitrogen and stored for later use.

### RNA production

tRNA precursors were prepared by T7 in-vitro transcription. DNA templates were synthesized (Integrated DNA technologies), amplified by PCR (Phusion High-Fidelity DNA polymerase kit, Thermo Fisher Scientific) and purified by phenol:chloroform extraction. Run-off transcription reactions were assembled with in-house purified T7 RNA polymerase (40 mM Tris pH 8, 20 mM MgCl_2_, 2 mM spermidine, 0.05% tween-20, 10 mM TCEP, 4 mM each nucleotide, 5 μg DNA template) and run for 4 h at 37 °C. The transcripts were purified in a Mono Q ion exchange column (Cytiva) in the buffer 20 mM HEPES pH 7.5, 1 mM MgCl_2_ and 100 mM NaCl with a gradient up to 1 M NaCl.

### Mitochondrial RNAse Z complex reconstitution

TRMT10C/SDR5C1 was run over an S200 10/300 column (Cytiva) in RNAse-free buffer D (20 mM HEPES 7.5, 50 mM NaCl, 1 mM MgCl_2_, 1 mM DTT). ELAC2 was purified similarly but in 150 mM NaCl. The complex was assembled at 4°C with 10 μM TRMT10C/SDR5C1, 20 μM ELAC2 and 20 μM mt-tRNA precursor in buffer D with 200 μM S-(5’-Adenosyl)-L-methionine (SAM) (Sigma-Aldrich). The RNA and ELAC2 contained a higher NaCl concentration; therefore, the salt concentration was adjusted to 50 mM NaCl using the same buffer without NaCl. The complex was applied over the same S200 size exclusion chromatography in buffer D. RNAse Z was concentrated by ultrafiltration, supplemented with 0.005% Tween-20 and 200 μM SAM, and directly used for grid preparation.

### RNAse Z cleavage assays

The assays were performed as previously described (Reinhard et al., 2017). Briefly, the reaction was assembled in a buffer containing 20 mM HEPES/KOH pH 7.6, 130 mM KCl, 2 mM MgCl_2_, 2 mM TCEP, 5 μM SAM, and 0.1 mg/ml bovine serum albumin (BSA). The RNA substrates (400 nM) were pre-incubated for 10 min at room temperature with TRMT10C/SDR5C1 (1600 nM). Then, they were mixed 1:1 with ELAC2 (800 nM to 50 nM in a 1:2 dilution series). The reactions were incubated for 20 min at room temperature. Proteinase K (RNAse grade, Thermo Fisher Scientific) was added to each reaction to a final concentration of 0.4 mg/ml and further incubated at 50 °C for 10 min. The reactions were analyzed using 15 % TBE-UREA PAGE (Thermo Fisher Scientific) and stained with SYBR Green II RNA stain (Thermo Fisher Scientific).

### Grid preparation and data collection

UltrAuFoil 300 mesh (Quantifoil Micro Tools GMBH; R 1.2/1.3 geometry) grids were glow-discharged at 25 mA for 30 s using an EMS100X glow-discharge unit. Four μl of sample with an OD_260_ of 7.5 were applied to the grids and vitrified at 4 °C and 100 % humidity using a Vitrobot Mk IV (Thermo Fisher Scientific) (blot 6s, blot force 3, 595 filter paper (Ted Pella Inc.)).

All cryo-EM data collection was performed with EPU (Thermo Fisher Scientific) using a Krios G3i transmission electron microscope (Thermo Fisher Scientific) operated at 300kV in the Karolinska Institutet’s 3D-EM facility. Images were acquired in nanoprobe 165kX EF-TEM SA mode (0.505 Å/px) with a slit width of 10eV using a K3 Bioquantum, for 1.5s with 60 fractions and a total fluency of 48 e^-^/Å^2^. Motion correction, dose weighting, CTF estimation, Fourier cropping (to 1.01Å/px), particle picking (size threshold 100Å, using the pre-trained BoxNet2Mask_20180918 model), and extraction in 416-pixel boxes were performed on the fly using Warp (Tegunov & Cramer, 2019). Only particles from micrographs with an estimated resolution of 4 Å or better and under-focus between 0.2 and 3 μm were retained for further processing.

### Data processing and model building

The particles were further processed using CryoSPARC v4.1 (Punjani et al., 2017). The RNAse Z-HS particles were subjected to 2D classification. Particles from good and bad 2D classes were used to generate one and four ab initio volumes, respectively. The RNAse Z-HS and RNAse Z^H548A^-HS dataset were processed similarly. First, the particle sets were cleaned by heterogeneous refinement using the five ab initio volumes from above. Then, 3D-variability analysis (Punjani & Fleet, 2020) was performed to identify the ELAC2-containing particles. In the case of RNAse Z^H548A^-HS, two rounds of heterogeneous refinement were performed, and the 3D-variability analysis was replaced by a 3D classification, which fulfilled a similar function in selecting the ELAC2-containing particles. Then, a local refinement focused on ELAC2 was performed. Finally, a 3D classification with a resolution cutoff of 3 Å was performed for final dataset cleanup. One of the classes yielded a reconstruction with a resolution approximating 3 Å, while the other classes remained in the range of 4-5 Å. The same particle set was subjected to a non-uniform refinement with a mask covering the entire RNAse Z.

The ELAC2-focused local refinement and the non-uniform refinement were combined in Phenix (Liebschner et al., 2019). To ensure that ELAC2 is correctly positioned with respect to the rest of the particle, the ELAC2-focused map was fitted into the ELAC2 map from the non-uniform refinement before, combining the global and focused maps. For the RNAse Z-HCCA structure, the dataset was processed as follows. First, the particle set was cleaned by heterogeneous refinement. The ELAC2-containing particles were then selected by 3D classification using a focus mask on the ELAC2 region. These particles were then refined by 3D non-uniform refinement. The dataset was thereafter transferred to RELION-5.0 beta (Scheres, 2012). In RELION, 3D auto-refinement and CTF refinement (beamtilt, trefoil, anisotropic magnification and per-particle defocus) were performed. Next, particle subtraction was performed to focus on ELAC2. A 3D classification was performed with local pose searches and a regularization T of 20 to select the ELAC2 particles that refine to high resolution. A final round of per-particle defocus refinement and 3D auto-refine with Blush regularization (Kimanius et al., 2023) was performed. The consensus map and the ELAC2-focused map were combined in Phenix.

For atomic model building, a starting model for RNAse Z was generated using the RNAse P structure (Bhatta et al., 2021; PDB 7ONU) for TRMT10C, SDR5C1, and the mt-tRNA precursor, and AlphaFold (Jumper et al., 2021) for ELAC2. The model was built and refined using Coot (Emsley et al., 2010) and Phenix (Liebschner et al., 2019). For the low-resolution areas (the ELAC2 exosite), the model was refined using Isolde (Croll, 2018) instead.

The combined maps were improved iteratively by postprocessing with Locscale2 (Jakobi et al., 2017) and model building in Coot/Isolde.

## Data availability

Publicly available datasets from Protein Data Bank (7ONU, 4GCW, 1Y44) were used for atomic model building and comparison. Cryo-EM maps and atomic coordinates of the reported structures were deposited in Electron Microscopy Data Bank (EMDB) and Protein Data Bank, respectively, with the following accession codes: PDB-9EY0, EMD-50050, 19954 and 19955 (RNAse Z-HS); PDB-9EY1, EMD-50051, 19956 and 19957 (RNAse Z^H548A^-HS); PDB-9EY2, EMD-50052, 19958 and 19959 (RNAse Z-HCCA).

Source data are provided in this paper.

## Acknowledgments

The work was funded by the Knut & Alice Wallenberg Foundation (KAW 2017.0080 and 2018.0080 awarded to BMH) and the Swedish Research Council (VR 2018-03808 and 2022-02326 awarded to BMH). All cryo-EM data was collected at the Karolinska Institutet’s 3D-EM facility, and we thank A. Bondy for excellent support.

## Conflict of Interests

The authors declare that they have no conflict of interest.

## Notes

### Competing Interest Statement

The authors have declared no competing interest.

