## Supplemental Figures for "Structural basis of 3′-tRNA maturation by the human mitochondrial RNAse Z complex"


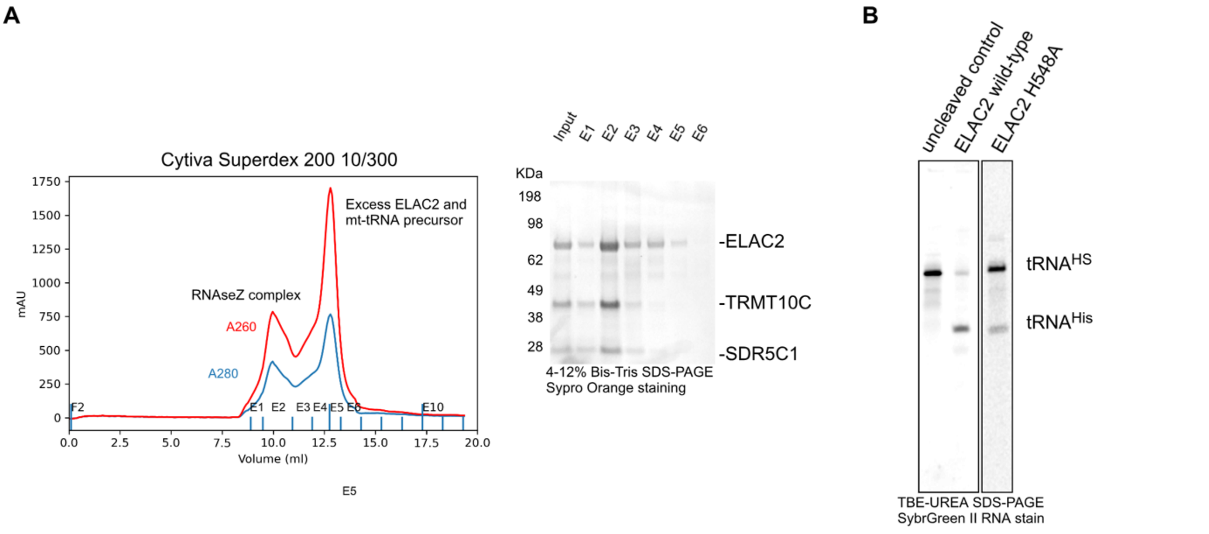


**Supplementary Figure 1. Mitochondrial RNAse Z complex formation. A.** TRMT10C/SDR5C1 was mixed with a twofold molar excess of ELAC2 and mt-tRNA-HS precursor and subjected to size exclusion chromatography. The fractions E1-E6 were analyzed by SDS-PAGE stained for protein (middle panel). Fraction E2 was used for grid preparation. **B**. TBE-UREA polyacrylamide gel electrophoresis of the samples used for grid preparation, with wild-type ELAC2 and the H548A mutant. Please note that the ELAC2-H548A mutant still cleaves its substrate but very slowly.

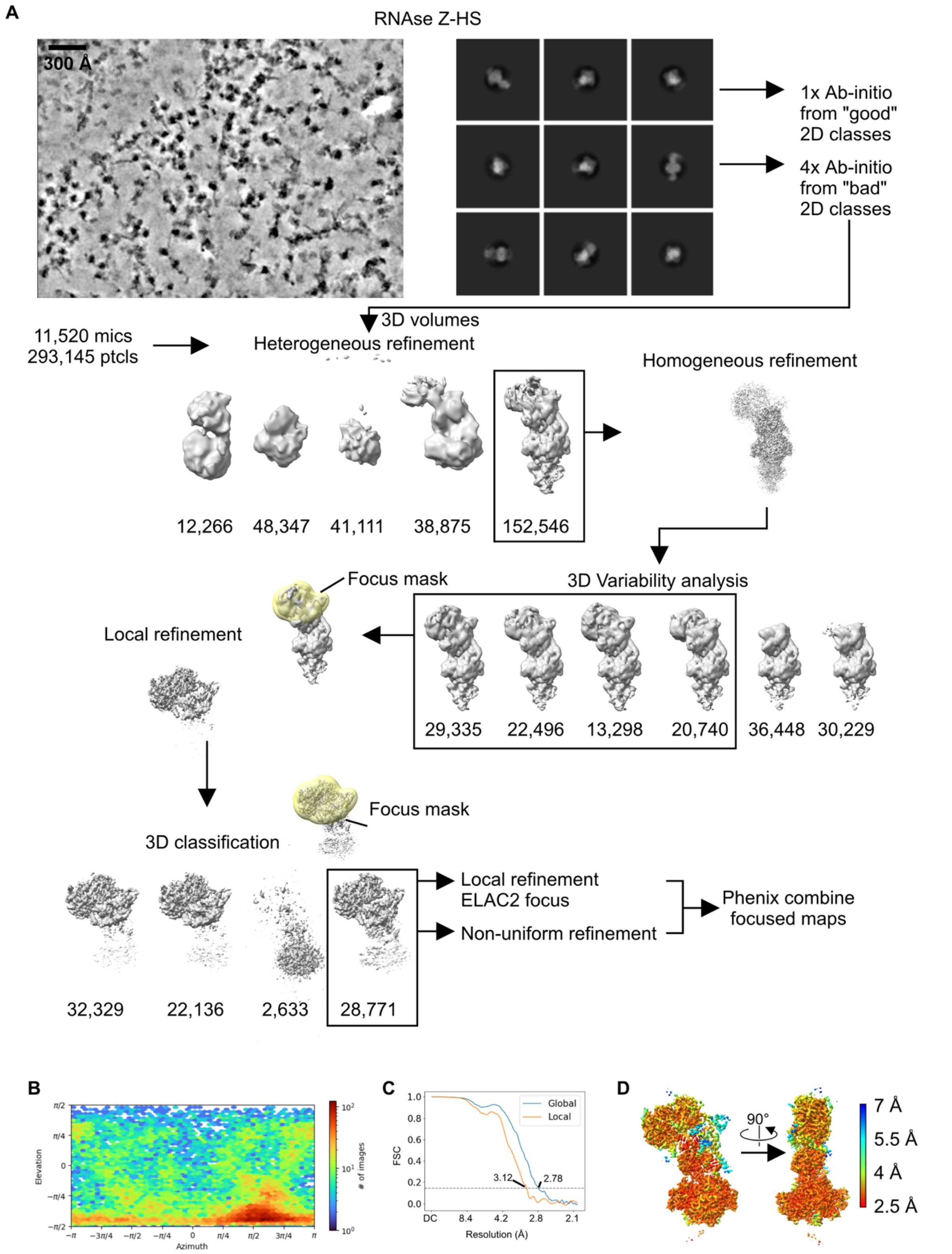


**Supplementary Figure 2. Cryo-EM data processing for the RNAse Z-HS dataset**. **A.** Data processing strategy. All the micrographs were preprocessed in WARP. An example micrograph denoised in WARP is shown. The particles extracted by WARP were processed using CryoSPARC v4.1. Examples of good 2D classes are shown. **B.** Angular orientations of the particles used in the final reconstruction. **C.** Fourier Shell Correlation of the final global and local refinements. **D.** Composite map colored by local resolution, as calculated in CryoSPARC using the composite half maps.

**A**

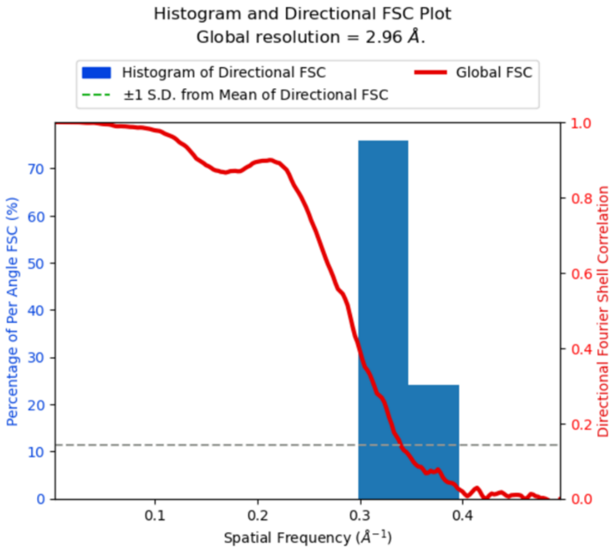


**B**

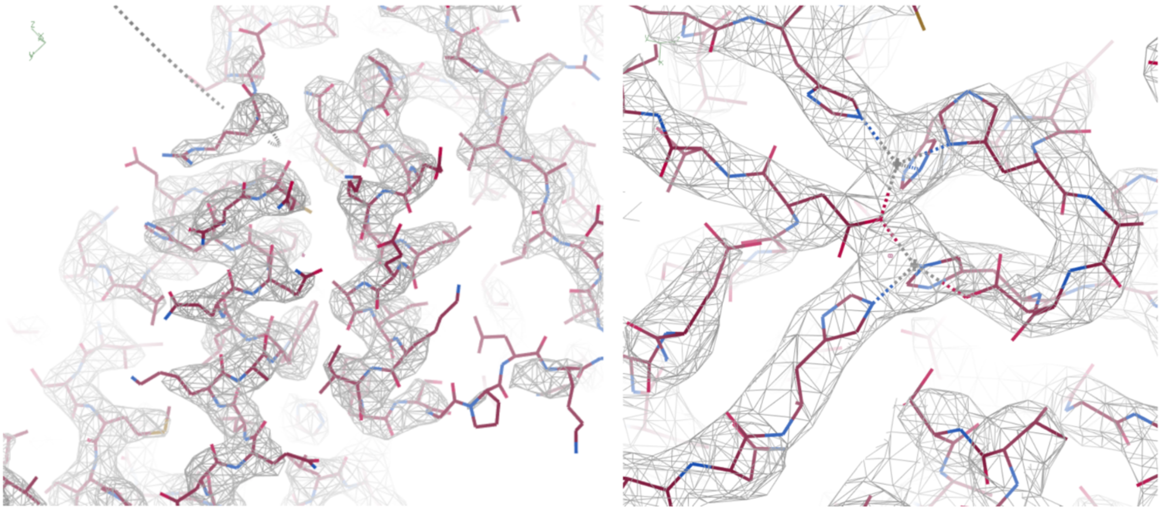

**Supplementary Figure 3. Resolution of the RNAse Z-HS cryo-EM structure. A.** Histograms and directional FSC plots for the EM density map calculated using 3DFSC (Tan et al., 2017). The composite half maps with an auto-tightened FSC mask from CryoSPARC were provided as input (cone angle 20 degrees, FSC cutoff 0.143, Sphericity threshold 0.5, and high pass filter 150 Å). **B**. Representative non-carved raw maps (RMSD 2.0).

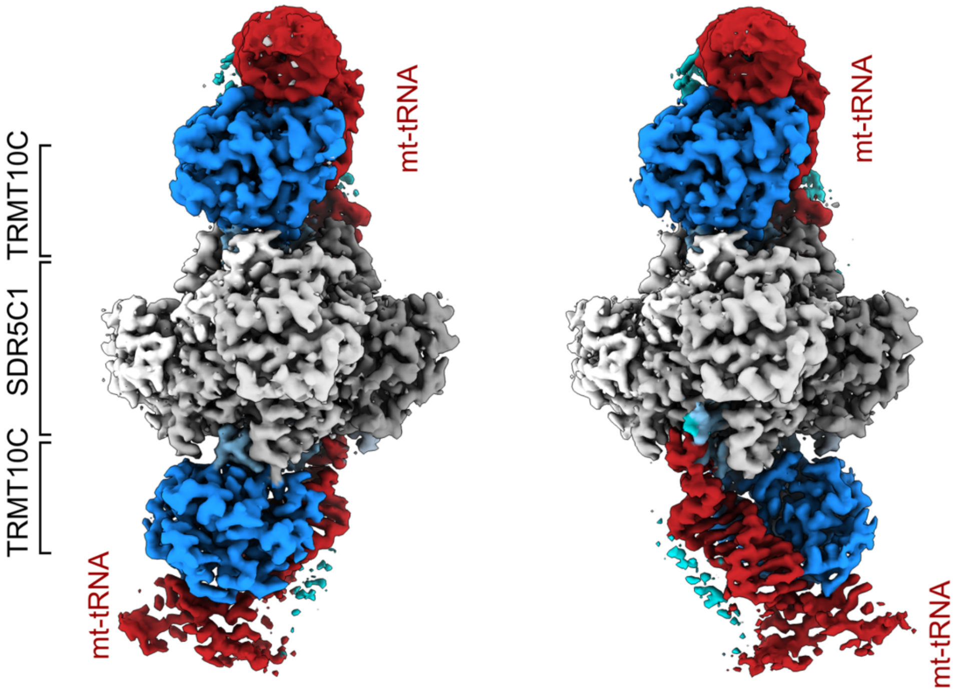


**Supplementary Figure 4. Pseudosymmetry in mitochondrial RNAse Z.** Cryo-EM density maps showing the two possible RNAse Z complexes. The 4xSDR5C1 platform offers two mt-tRNA/TRMT10C binding sites. The 4xSDR5C1 platform has a vertical and a horizontal two-fold symmetry axis. Consequently, it supports two mt-tRNA/TRMT10C subcomplexes that can be in two different orientations with respect to each other (left and right panels). The 4xSDR5C1 are colored in different shades of gray; TRMT10C is colored in dark blue with the NTD highlighted in light blue; the mt-tRNA is in red. The ELAC2 density is not visible due to its low occupancy in the particles used for these examples.

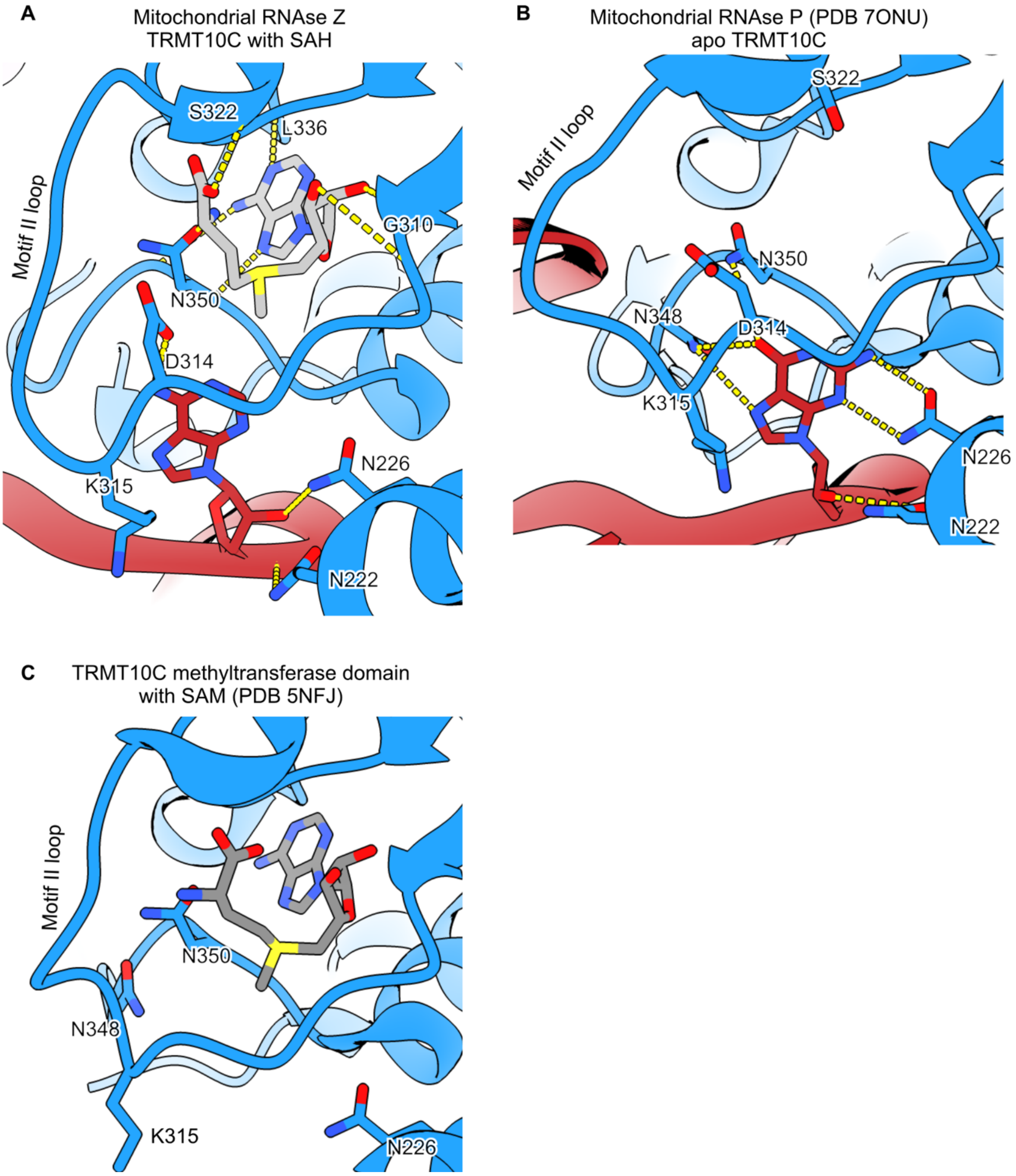


**Supplementary Figure 5. TRMT10C active site. A.** TRMT10C carries S-adenosyl homocysteine (SAH) in the active site. The mt-tRNA nucleotide at position 9 (adenine) carries the N1 methyl group. The electrostatic interactions established with SAH and A9 are indicated. **B and C.** Comparison with the apo TRMT10C (in the mitochondrial RNAse P structure) (B) and the SAM-bound TRMT10C methyltransferase domain crystal structure (PDB 5NFJ) (C). The motif II loop switches conformation between the three structures.

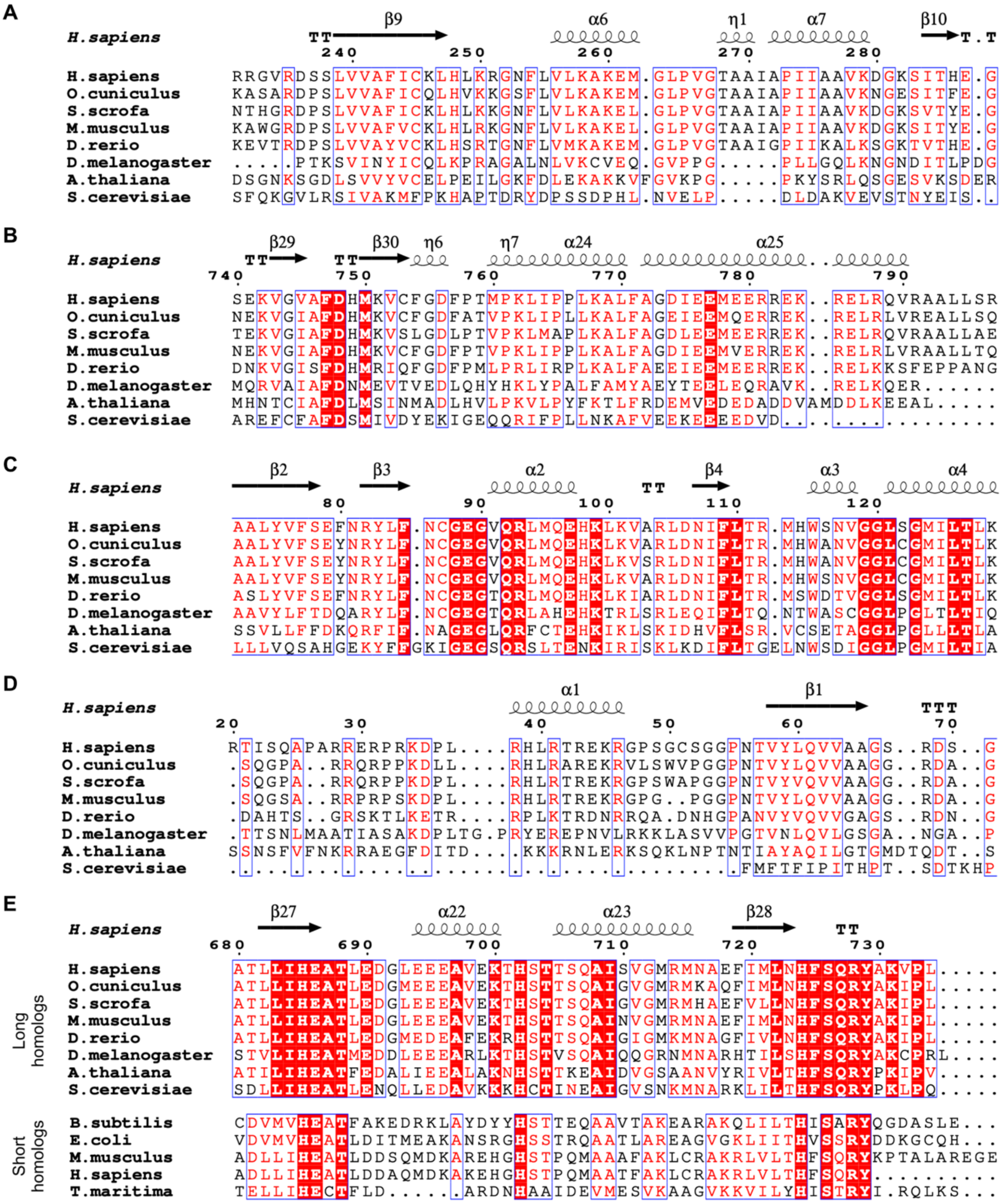


**Supplementary Figure 6. Structure-based multiple sequence alignment of ELAC2 homologs. A-E.** Multiple segments of the multiple sequence alignment. The alignment was generated using Promals3D (Pei et al., 2008). The UniProt IDs/RefSeq and PDB codes (when available) are as follows. Long homologs: *H. sapiens* ELAC2 (Q9BQ52, RNAse Z structure); *O. cuniculus* (XM_002718909.4); *S. scrofa* (NM_001243216.1); *M. musculus* ELAC2 (Q80Y81); *D. rerio* (NP_001243133.1); *D. melanogaster* (Q8MKW7); *A. thaliana* ELAC2 (Q8VYS2); *S. cerevisiae* ELAC2 (P36159, PDB 5MTZ). Short homologs: *B. subtilis* RNAse Z (P54548, PDB 1Y44); *E. coli* RNAseBN (P0A8V0); *M. musculus* ELAC1 (Q8VEB6); ELAC1 (Q9H777, PDB 3ZWF); and *T. maritima* (Q9WZW8, PDB 2E7Y).

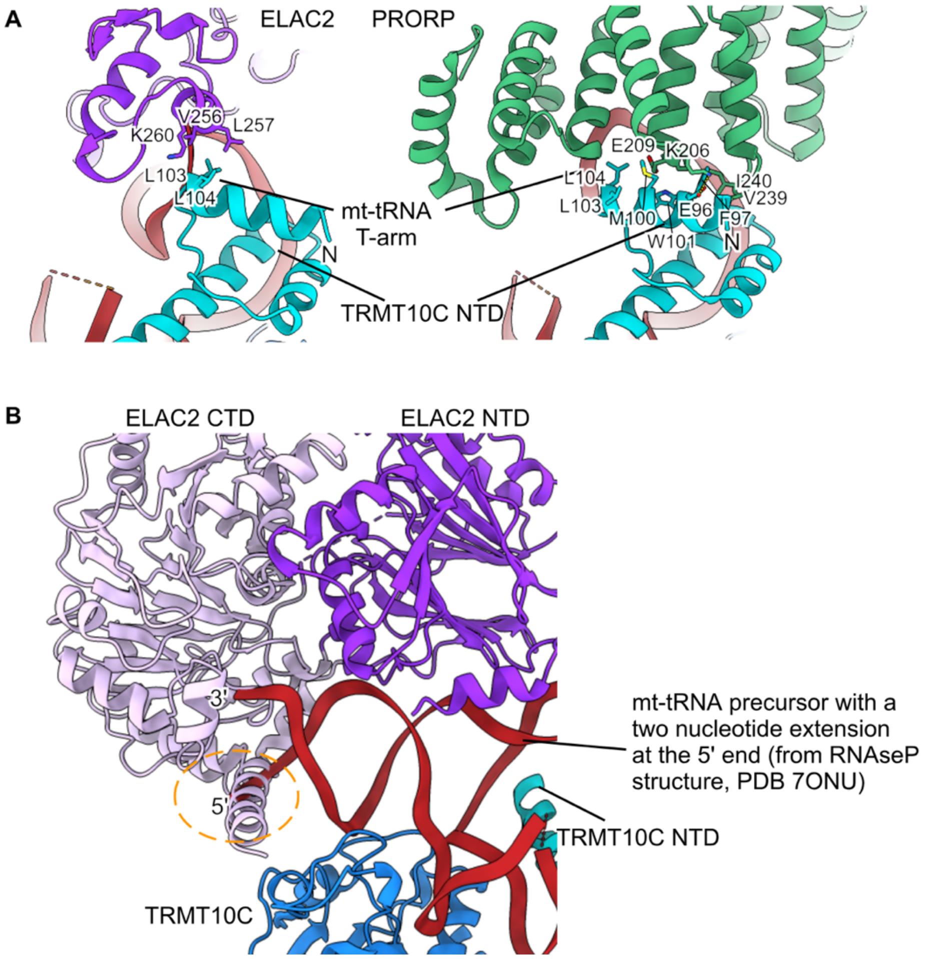


**Supplementary Figure 7. Comparison of RNAse Z and RNAse P. A.** TRMT10C NTD interacts with ELAC2 (RNAse Z, left panel) and PRORP (RNAse P, right panel), in the T-arm loop region. **B.** The ELAC2 C-terminal helix would collide with a 5' extension on the mt-tRNA precursor (yellow circle). To make this figure, the mt-tRNA precursor in RNAse Z was replaced with the RNAse P precursor (mt-tRNA$^{Tyr}$), which has a 2-nucleotide 5' extension. For this, The RNAse Z structure was superimposed on the RNAse P structure (Bhatta et al., 2021; PDB 7ONU), using TRMT10C as a reference.

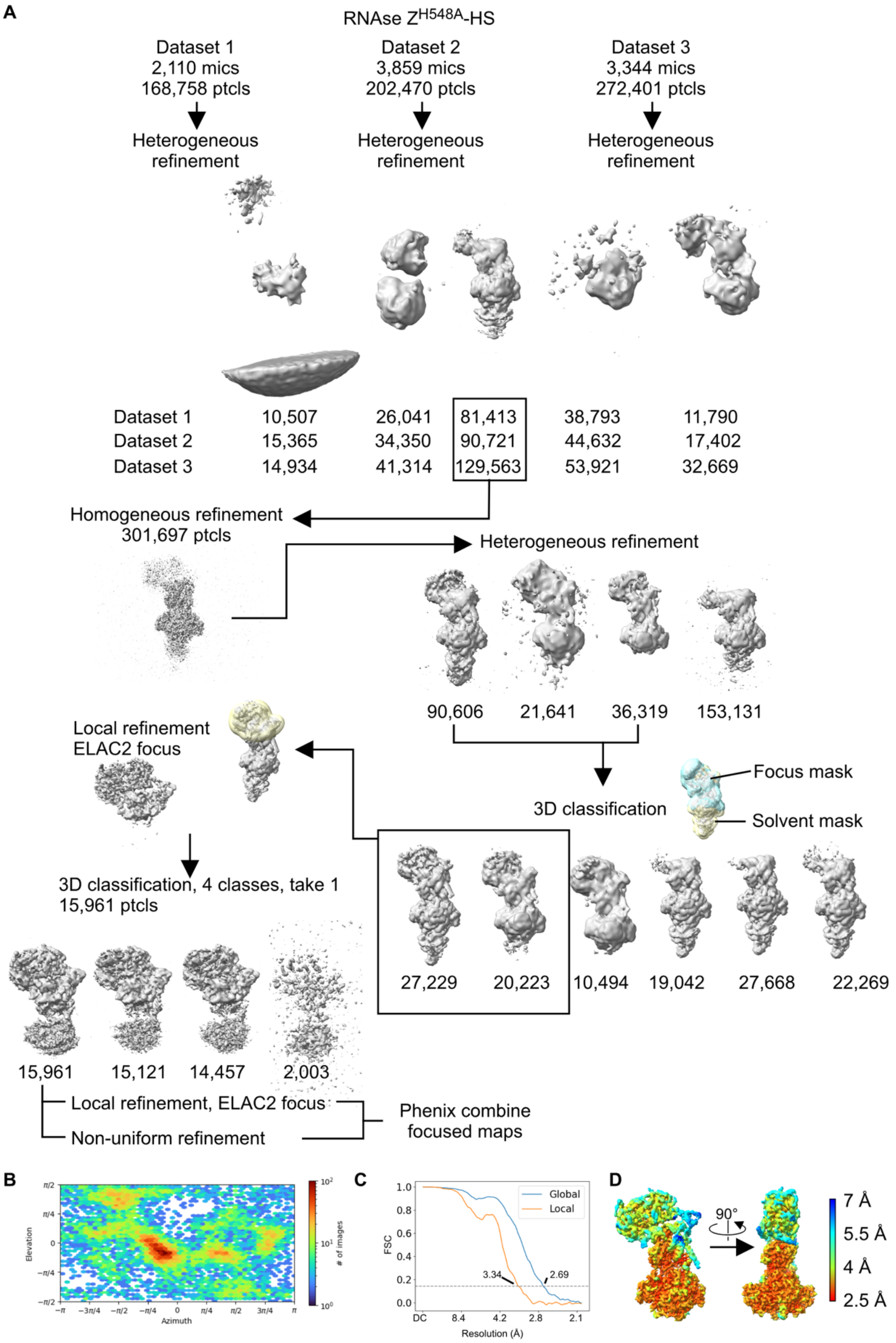


**Supplementary Figure 8. Cryo-EM data processing for the RNAse Z$^{H548A}$-HS dataset**. **A.** Data processing strategy. All the micrographs were preprocessed in WARP. Three datasets were pooled after independent heterogeneous refinement jobs, using the *ab initio* volumes from RNAse Z-HS (Supplementary Fig.2) as 3D references. **B.** Angular orientations of the particles used in the final reconstruction. **C.** Fourier Shell Correlation of the final global and local refinements. **D.** Composite map colored by local resolution, as calculated in CryoSPARC using the composite half maps.

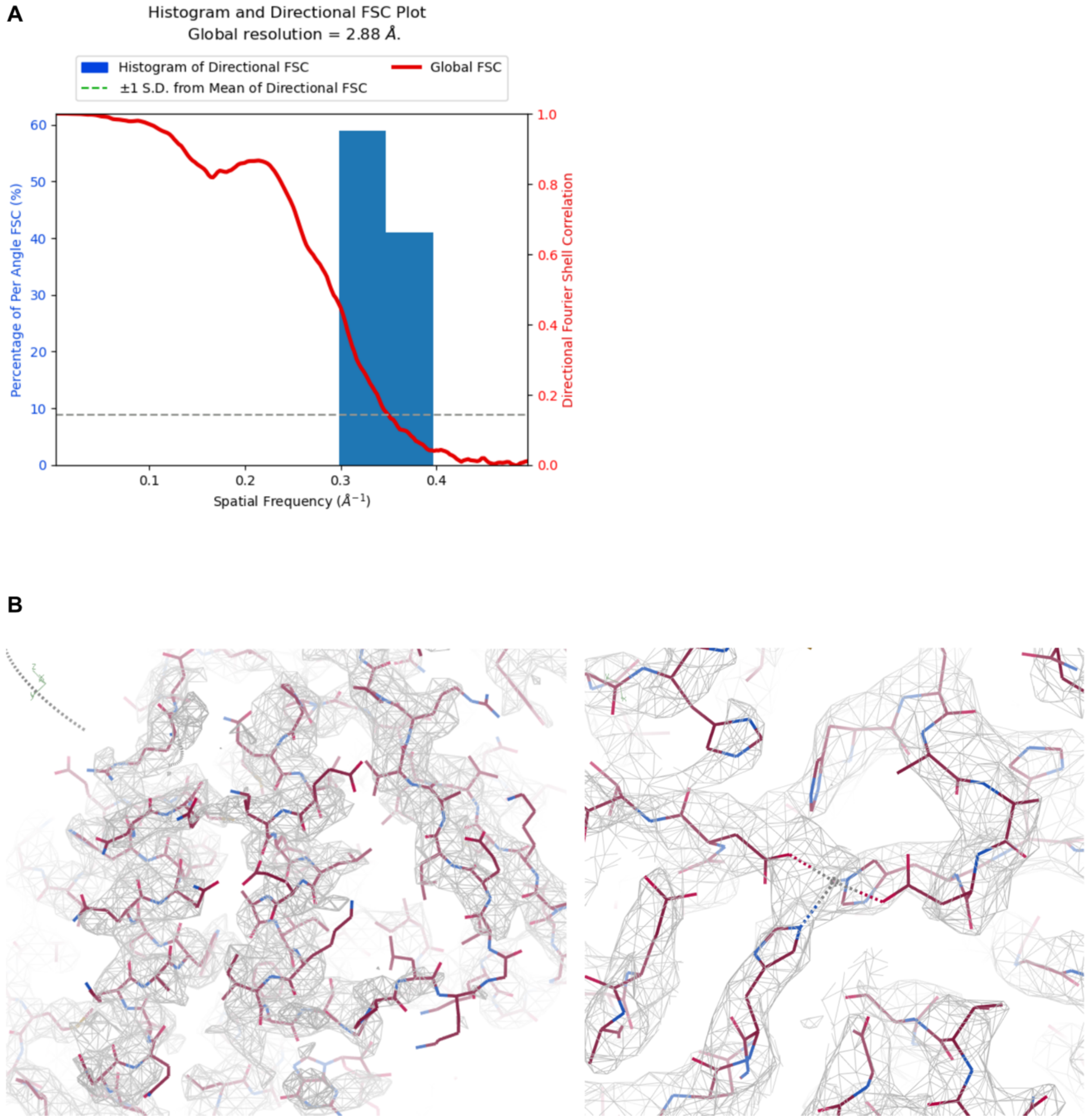


**Supplementary Figure 9. Resolution of the RNAse Z$^{H548A}$-HS cryo-EM structure. A.** Histograms and directional FSC plots for the EM density map calculated using 3DFSC (Tan et al., 2017). The composite half maps with a auto-tightened FSC mask from CryoSPARC were provided as input (cone angle 20 degrees, FSC cutoff 0.143, Sphericity threshold 0.5, and high pass filter 150 Å). **B**. Representative non-carved raw maps (RMSD 2.0).

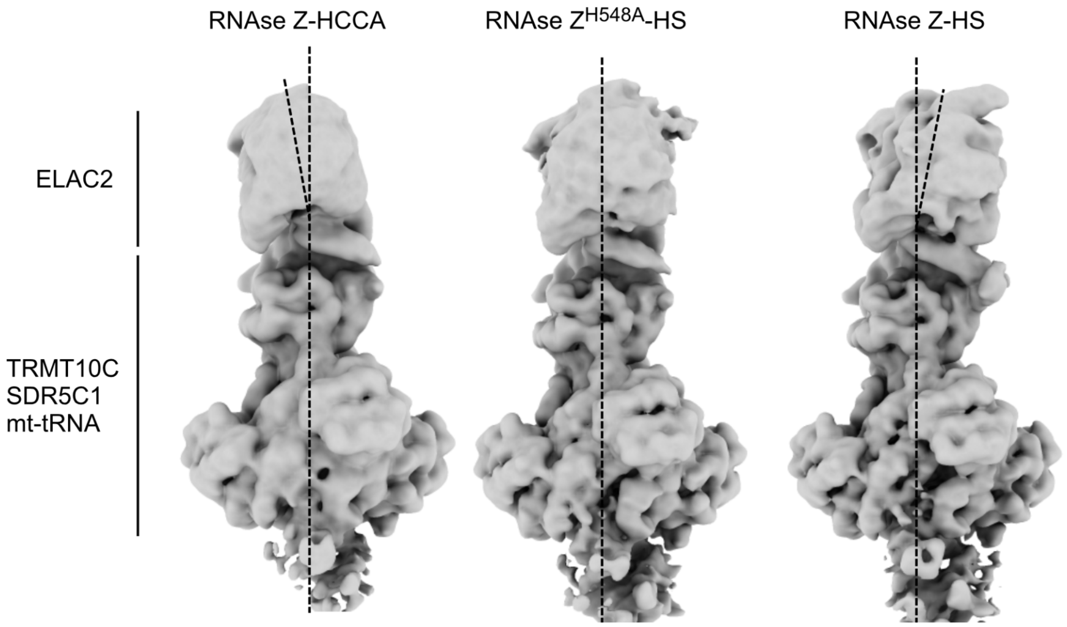


**Supplementary Figure 10. ELAC2 pivots on the mt-tRNA.** The consensus refinement maps are shown. ELAC2 corresponds to the top part of the map, whereas TRMT10C, SDR5C1, and mt-tRNA correspond to the bottom part. A long vertical dotted line represents the main vertical axis of the particle, whereas the short line represents the main axis of ELAC2. The intersection between the two represents the hinge point.

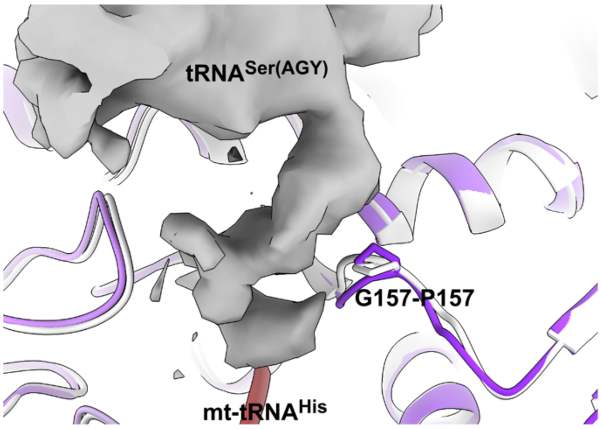


**Supplementary Figure 11. Cryo-EM map for mt-tRNA$^{Ser(AGY)}$ in the RNAse Z$^{H548A}$-HS structure.** The density, for which it was not possible to create an atomic model, displaces the G157-P157 loop compared with the RNAse Z-HS structure (superposed in white)

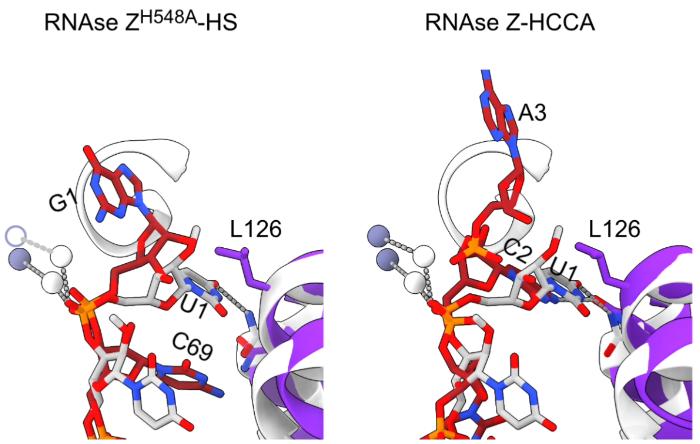


**Supplementary Figure 12. Comparison with the *B. subtilis* RNAse Z bound to precursor tRNA.** In *B. Subtilis* (white), U1 inserts into a pocket and helps position the scissile bond. In RNAse Z$^{H548A}$-HS, L126 partially blocks the pocket for G1; nevertheless, the backbone remains the same. In RNAse Z-HCCA (right panel), U1 corresponds very well to C2, but the backbone is distorted so that the 3′-CCA tail is not cleaved. The *B. Subtilis* $Zn^{2+}$ are shown as white spheres. The correspondence with the human structures is shown with gray dotted lines. The $Zn^{2+}$ are further away in ELAC2 as the active site channel is wider. In RNAse Z$^{H548A}$-HS, the missing $Zn^{2+}$ is drawn based on the RNAse Z-HS structure.

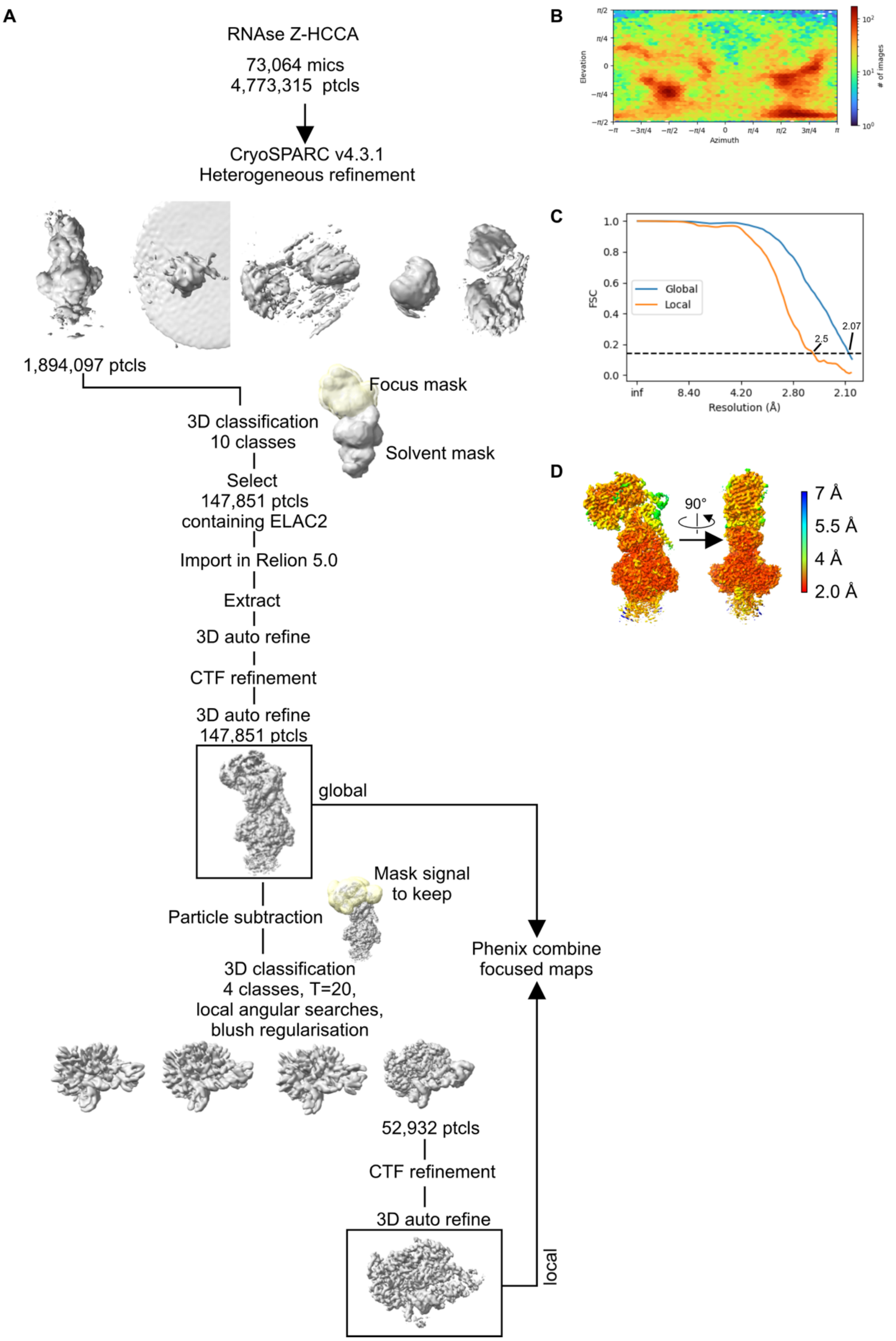


**Supplementary Figure 13. Cryo-EM data processing for the RNAse Z-HCCA dataset**. **A.** Data processing strategy. All micrographs were pre-processed in WARP. The particles extracted by WARP were processed in CryoSPARC v4.1 before re-extraction in RELION for further processing **B.** Angular orientations of the particles used in the final reconstruction. **C.** Fourier Shell Correlation of the final global and local refinements. **D.** Composite map colored by local resolution, as calculated in CryoSPARC using the composite half maps.

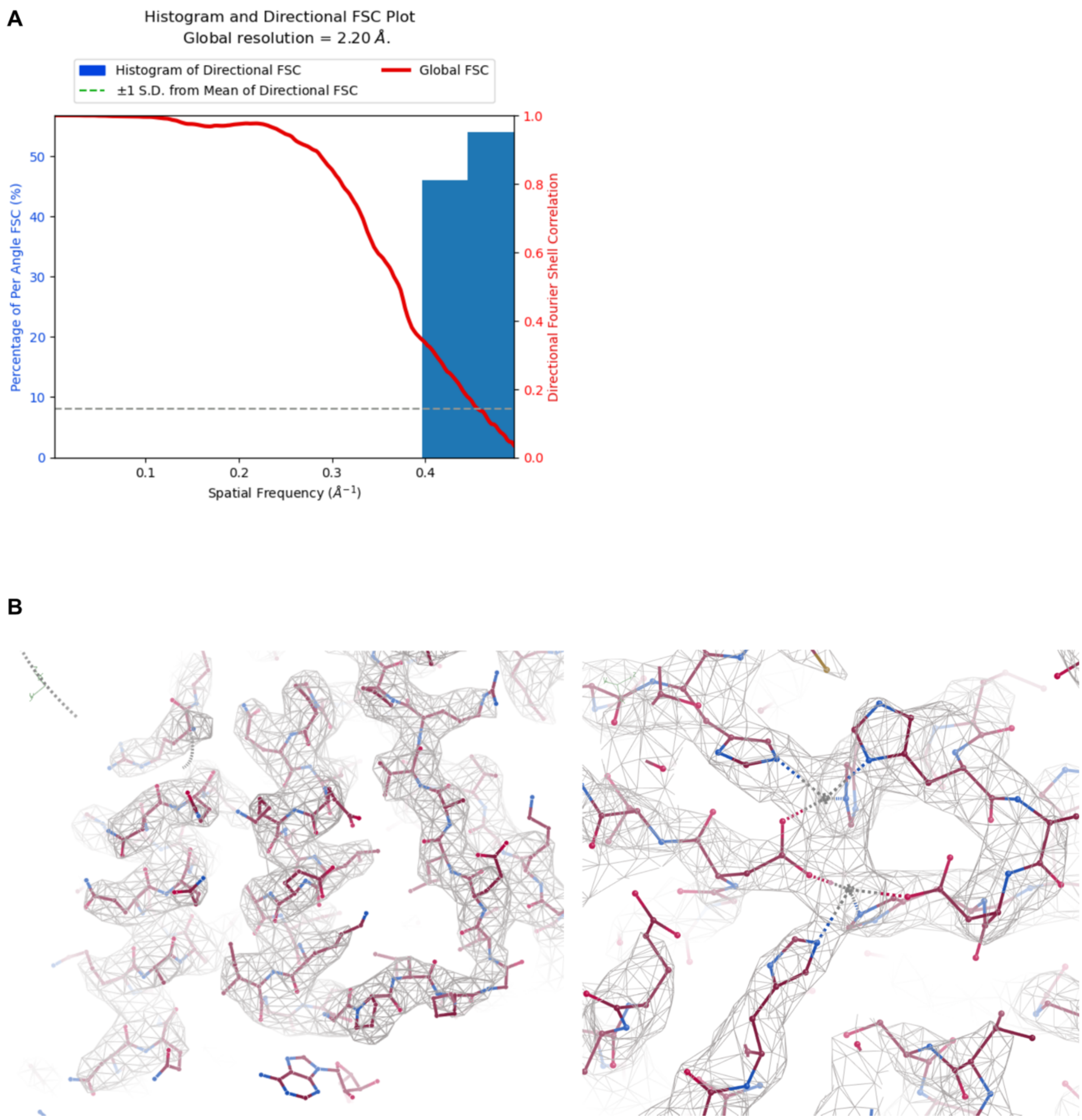


**Supplementary Figure 14. Resolution of the RNAse Z-HCCA cryo-EM structure. A.** Histograms and directional FSC plots for the EM density map calculated using 3DFSC (Tan et al., 2017). The composite half maps with an auto-tightened FSC mask from CryoSPARC were provided as input (cone angle 20 degrees, FSC cutoff 0.143, Sphericity threshold 0.5, and high pass filter 150 Å). **B**. Representative non-carved raw maps (RMSD 2.0).
